# Integrative analysis of scRNAs-seq and scATAC-seq revealed transit-amplifying thymic epithelial cells expressing autoimmune regulator

**DOI:** 10.1101/2021.10.04.463004

**Authors:** Takahisa Miyao, Maki Miyauchi, S. Thomas Kelly, Tommy W. Terooatea, Tatsuya Ishikawa, Eugene Oh, Sotaro Hirai, Kenta Horie, Yuki Takakura, Houko Ohki, Mio Hayama, Yuya Maruyama, Takao Seki, Haruka Yabukami, Masaki Yoshida, Azusa Inoue, Asako Sakaue-Sawano, Atsushi Miyawaki, Masafumi Muratani, Aki Minoda, Nobuko Akiyama, Taishin Akiyama

## Abstract

Medullary thymic epithelial cells (mTECs) are critical for self-tolerance induction in T cells via promiscuous expression of tissue-specific antigens (TSAs), which are controlled by transcriptional regulator AIRE. Whereas AIRE-expressing (Aire^+^) mTECs undergo constant turnover in the adult thymus, mechanisms underlying differentiation of postnatal mTECs remain to be discovered. Integrative analysis of single-cell assays for transposase accessible chromatin (scATAC-seq) and single-cell RNA sequencing (scRNA-seq) suggested the presence of proliferating mTECs with a specific chromatin structure, which express high levels of Aire and co-stimulatory molecules CD80 (Aire^+^CD80^hi^). Proliferating Aire^+^CD80^hi^ mTECs detected by using Fucci technology express a minimal level of Aire-dependent TSAs and are converted into quiescent Aire^+^CD80^hi^ mTECs expressing high levels of TSAs after a transit amplification. These data provide evidence for the existence of transit amplifying Aire^+^mTEC precursors during Aire^+^mTEC differentiation process of the postnatal thymus.

## Introduction

Medullary thymic epithelial cells (mTECs) are essential for induction of T cell self-tolerance in the thymus^1, 2^. mTECs ectopically express thousands of tissue-specific antigens (TSAs), and this expression is regulated by transcription factors, AIRE and FEZF2^3, 4^. TSAs are directly or indirectly presented to developing T cells, and T cells that recognize TSAs with high affinity undergo apoptosis or are converted into regulatory T cells, thereby suppressing the onset of autoimmune diseases^1, 2^.

Several studies have suggested processes and underlying mechanisms of mTEC differentiation during thymic organogenesis^1, 2, 5, 6, 7, 8, 9, 10, 11^. In addition, some previous studies suggest that mTEC turnover is homeostatic in the adult thymus, with a duration of approximately 2 weeks^12, 13, 14^. Notably however, cellular mechanisms underlying maintenance of adult mTECs remain unclear. mTEC subpopulations are largely classified based on their expression of cell surface markers (mainly CD80 and MHC class II) and Aire in the adult thymus^1^. CD80^lo^ and Aire-negative (Aire^-^) mTECs (mTEC^lo^) are thought to be immature, and they differentiate into CD80^hi^ Aire-expressing (Aire^+^) mTECs that are reportedly post-mitotic^13^. Aire^+^ mTECs are further converted into Aire-negative mTECs (post-Aire mTECs)^15, 16, 17, 18, 19^. Moreover, a previous study suggested that mTECs might be differentiated from stage-specific embryonic antigen-1^+^ (SSEA-1) claudine3/4^+^ mTEC stem cells^20^. These views are primarily based on fate mapping studies involving transfer and re-aggregation of sorted cell populations with fetal thymus^5, 13, 20^ and on experiments employing genetic marking^15, 17^.

Single-cell RNA sequencing (scRNA-seq) technology has yielded new insights into cell diversity and differentiation in various tissues. In TEC biology, previous scRNA-seq studies revealed a stochastic nature of TSA expression in mTECs^21, 22^ and high heterogeneity of TECs in mice^23, 24, 25, 26^. Bornstein et al. showed that mTECs in the postnatal thymus are separated into four subsets, mTEC I to IV^23^. In addition to the classical mTEC^lo^ (mTEC I), Aire^+^ mTEC (mTEC II), and post-Aire mTEC (mTEC III) types, a tuft-like mTEC subset (mTEC IV) was identified^23, 24^. Subsequent scRNA-seq studies suggested further heterogeneity of TECs, such as cilium TECs^25^, GP2^+^ TECs^25^, intertypical TECs^26^, neural TECs^26^, and structural TECs^26^, according to specific gene expression profiles. However, it has not yet been clarified whether this heterogeneity identified from gene expression profiles is correlated with differences in chromatin structure.

In general, transit-amplifying cells (TACs) are a proliferating cell population linking stem cells and differentiated cells^27^. TACs are short-lived and undergo differentiation after a few cell divisions. To date, the presence of TACs has been confirmed in some tissues such as intestines^28^, hair follicles^29^, and neurons^30^. Previous analyses of scRNA-seq data of murine adult TECs revealed a cell cluster expressing an abundance of cell-cycle regulated genes, which implies the presence of TACs for TECs (TA-TECs)^25, 31^. Computational trajectory analysis of scRNA-seq data suggested that this population might give rise to Aire-expressing mTECs^25, 26^. Intriguingly, another trajectory study predicted that this cell cluster might differentiate into Aire-expressing mTECs and an mTEC population expressing CCL21a^31^. However, because the TA-TEC candidate has not been isolated and specific marker genes of TA-TECs have not been reported, exact properties of TA-TECs, in addition to their cellular fates, remain to be clarified.

In this study, droplet-based scRNA-seq and scATAC-seq of murine TECs were performed to characterize TEC heterogeneity and differentiation dynamics. Integrative analysis of these data showed that Aire^+^ mTECs are separated into at least 2 clusters with different gene expression profiles and chromatin accessibilities. One of these Aire^+^ mTEC clusters exhibited high expression of cell cycle-related genes, which accords with a previously proposed TAC population of mTECs^25, 31^. By using the Fucci technology^32^, proliferating mTECs expressing Aire and maturation marker CD80 were isolated as TA-TEC candidates. This proliferating Aire^+^ CD80^hi^ mTEC subpopulation showed minimal expression of TSAs regulated by AIRE, in contrast to quiescent Aire^+^ CD80^hi^ mTECs. Moreover, *in vivo* BrdU pulse-labeling, and *in vitro* reaggregated thymic organ culture suggested that proliferating Aire^+^ CD80^hi^ mTECs are short-lived and that they differentiate into quiescent Aire^+^ CD80^hi^ mTECs, post-Aire mTECs, and tuft-like mTECs. Consequently, these data strongly suggest that proliferating Aire^+^ CD80^hi^ mTECs are TACs for mTECs expressing TSAs.

## Results

### Droplet-based scATAC-seq reveals heterogeneity of TEC chromatin structure

Given that chromatin structures can be changed during cell differentiation, scATAC-seq analysis of TECs may offer some insights into TEC heterogeneity and differentiation dynamics. Droplet-based scATAC-seq analysis was carried out with EpCAM^+^ CD45^-^ cells that were sorted and pooled from thymi of 2 mice, 4 weeks of age. Unsupervised graph-based clustering and dimensional reduction via uniform manifold approximation and production (UMAP) using the Signac R package (https://www.biorxiv.org/content/10.1101/2020.11.09.373613v1) revealed 11 cell clusters from 8,413 cells (Figure 1A). We first analyzed chromatin accessibility of previously known TEC marker genes. Clusters 0, 3, 4, 5, 8 and 9 contained relatively higher numbers of cells having the open chromatin structure of the *Cd80* gene, a maturation marker of TECs (Figure 1B and C). Among these clusters, the *cis*-regulatory element of the Aire gene^33^ (about 3 kbp upstream of the transcriptional start site) is opened in clusters 0 and 3 (Figure 1D), suggesting that these clusters (cluster 0 and 3) may be concordant with Aire-expressing mTECs (Aire^+^ mTECs, also referred to as mTEC II^23^). In contrast, the *cis* element of Aire genes is closed in clusters 4, 5, 8, and 9 (Figure 1D), suggesting that these clusters may correspond to post-Aire mTECs and other Aire-negative mature mTECs^23^. Because the *Lrmp* gene region is accessible in cluster 5 (Figure 1B and Figure 1—figure supplement 1A), this cluster may be equivalent to tuft-like mTECs (mTEC IV)^23, 24^. CD80 and Aire gene regions in clusters 1, 2, and 6 are relatively closed, whereas the mTEC marker *Tnfrsf11a* is relatively accessible (Figure 1B and C, and Figure 1—figure supplement 1A). Therefore, these clusters should be equivalent to mTECs expressing low levels of CD80 and Aire (mTEC^lo^). Cluster 7 should be cTECs, because cTEC marker *Psmb11* gene region is opened (Figure 1B and Figure 1—figure supplement 1A). Finally, cluster 10 was deemed thymocyte contamination because the *Rag1* gene was opened (Figure 1 —figure supplement 1A).

**Figure 1.**
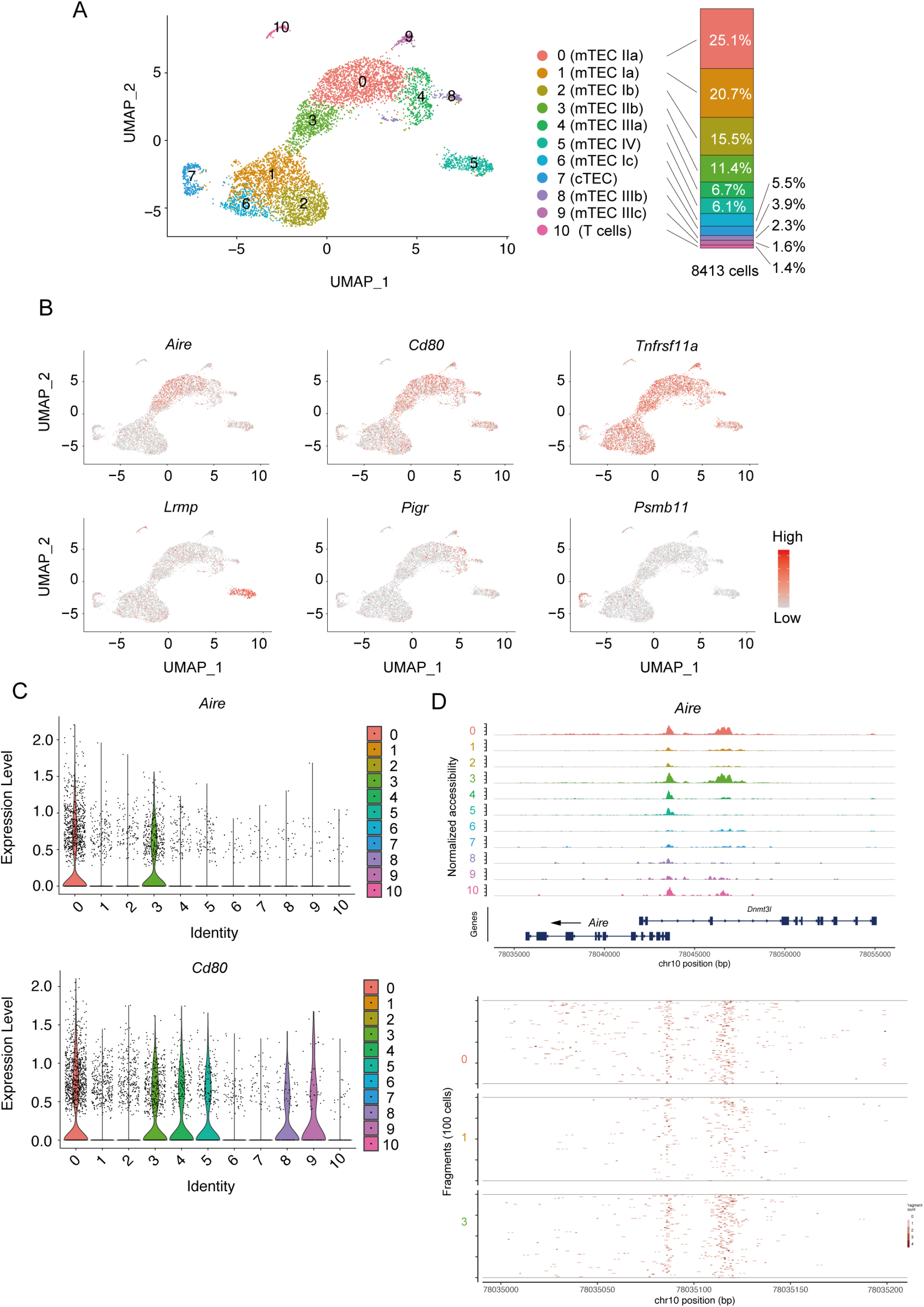
Droplet-based scATAC-seq analysis of TECs in 4-week-old mice. **A**. UMAP plot of scATAC-seq data from TEC cells (EpCAM^+^ CD45^-^ TER119^-^) from 4-week-old mice. Cell clusters are separated by colors and numbers in the plot. The graph on the right shows percentages of each cluster in the total number of cells detected (8413 cells). **B**. Chromatin accessibility of typical marker genes of TECs. Accessibility in each gene region is represented in red. **C.** Violin plot depicting chromatin accessibility in *Aire* and *Cd80* gene regions in each cluster. **D**. Pseudo-bulk accessibility tracks of the *Aire* gene region in each cluster (upper panels) and frequency of sequenced fragments within the *Aire* gene region of individual cells in cluster 0, 1 and 2 (lower panels)

We next sought to correlate the scATAC data with TEC scRNA-seq data. Droplet-based scRNA-seq analysis of 11,475 EpCAM^+^ CD45^-^ cells from age- and gender-matched mice (4-week-old female mice) revealed 18 cell clusters (Figure 2A), and expression of TEC marker genes in these clusters was analyzed (Figure 2B and Figure 2—figure supplement 1). Integrative analysis with reported datasets suggests quality comparable to that of our dataset (Figure 2—figure supplement 2). Clusters R0, R1, R3, and R9 showed high expression of *Aire*, suggesting that these clusters are equivalent to Aire^+^ mTECs (also referred to as mTECs II). Clusters R2, R4 and R5 include cells showing relatively higher levels of *Itga6* and *Ccl21a* expression and a very low level of *Aire* expression (Figure 2A and b), corresponding to mTEC I^23^, CCL21-expressing mTECs^34^, and possibly intertypical TECs^26^. Cluster R6 expresses *Lrmp* (Figure 2B), and should contain tuft-like mTECs (mTEC IV)^23^. Clusters R7 and R10 were marked with *Krt10* and *Pigr* genes, respectively (Figure 2B). Accordingly, these clusters should be categorized as post-Aire mTECs (also referred as to mTECs III^23^). Cluster R13 showed high expression of chemokines, *Ccl6* and *Gp2* (Figure 2B and Figure 2—figure supplement 1A), which should be concordant with Gp2^+^ TECs, as reported recently^25^. Clusters R8 and R11 exhibited high expression of typical cTEC marker genes, *Psmb11* and *Dll4* (Figure 2B and Figure 2—figure supplement 1A), and should be equivalent to cTECs. Given that thymocyte genes are highly expressed, cluster R11 was most likely thymic nurse cells enclosing thymocytes^35^. Cluster R12 showed relatively high expression of *Pdpn* (Figure 2—figure supplement 1A), which may comprise junctional TECs^36^. Cluster R14 was considered thymocyte contamination because thymocyte markers, but not TEC markers, were detected. Cluster R15 apparently corresponds to structural TECs, reported recently, because of their expression of *Car8* and *Cd177*^26^ (Figure 2—figure supplement 1A). Cells in cluster R16 highly express *Tppp3* and *Fam183b* (Figure 2—figure supplement 1A). Since these genes are expressed in ciliated cells^37, 38^, this cluster may be equivalent to ciliated columnar TECs associated with thymic cystic structure^25, 39, 40^. We failed to assign cluster R17, which may be contaminated with endothelial cells or macrophages, because they express *Ly6c1* and *Aqp1*, but low levels of *Epcam* (Figure 2—figure supplement 1A). Overall, our data and assignments are reasonably correlated with previous scRNA-seq data analyses^23, 25, 26, 31^.

**Figure 2.**
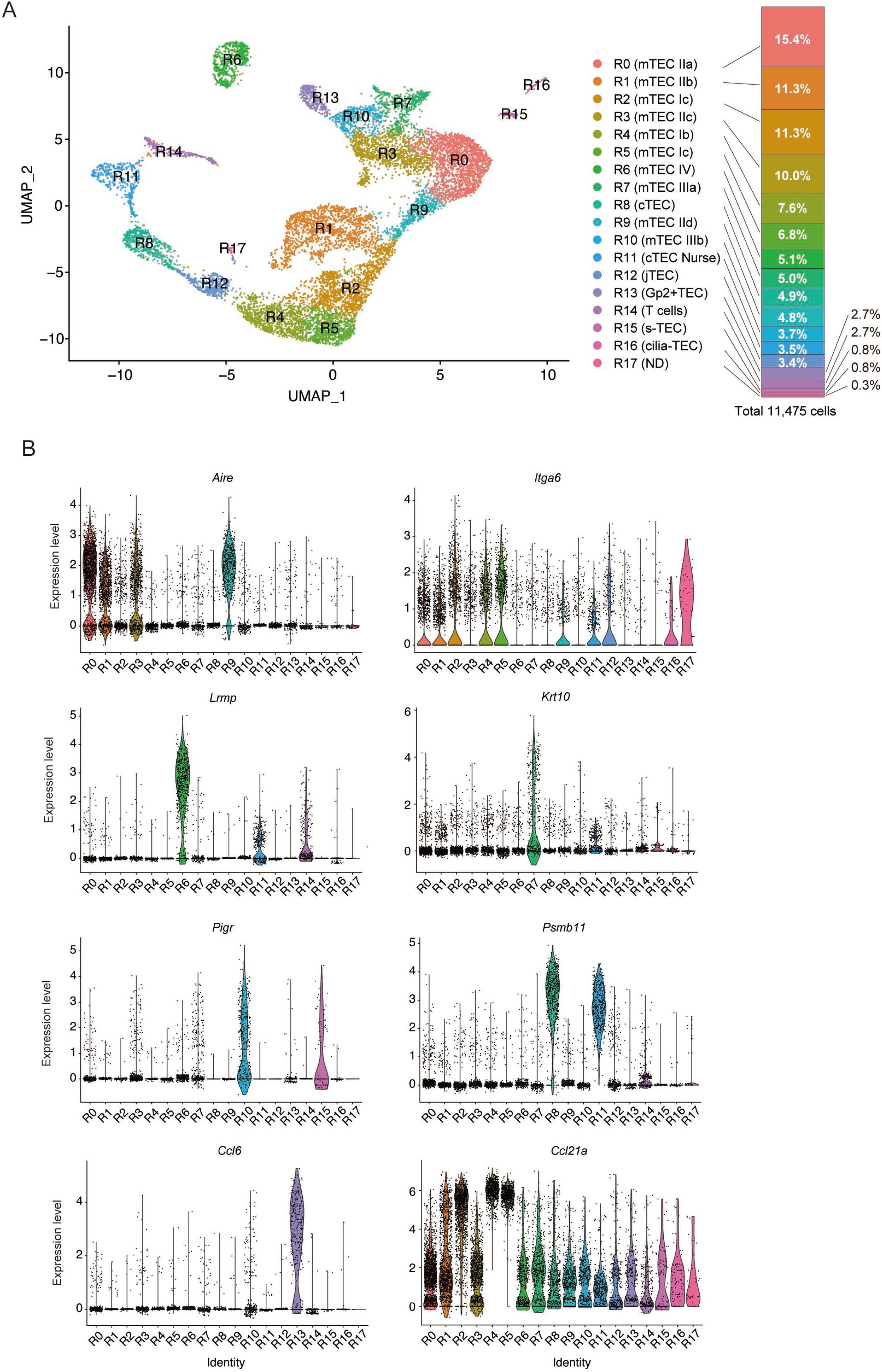
Droplet-based scRNA-seq analysis of TECs in 4-week-old mice. **A**. UMAP plot of scRNA-seq data from TEC cells (EpCAM^+^ CD45^-^ TER119^-^) from 4-week-old mice. Cell clusters (R0 to R17) are indicated by colors and numbers in the plot. The graph on the right shows the percentages of each cluster in the total number of cells detected (11,792 cells). **B**. Violin plots depicting expression level of typical TEC marker genes in each cluster.

We then bioinformatically integrated the scRNA-seq data with scATAC-seq data. Gene expression, predicted from accessible chromatin regions of scATAC-seq data, was correlated with scRNA-seq data using the Signac R package (Figure 3A and Figure 3—figure supplement 1). As described, clusters 0 and 3 in scATAC-seq analysis contain cells with the accessible *cis*-regulatory element of the *Aire* gene (Figure 1D). Consistently, cluster 0 in scATAC-seq were mostly transferred to cluster R0 (40.5 %) and R3 (26.3%) in scRNA-seq analysis (Figure 3B and C, and Supplementary Table 1), which were assigned as Aire^+^ mTECs (Figure 2). Cells transferred to R0 and R3 appear to be separately embedded in cluster 0 in the UMAP dimension, implying that these two Aire^+^ mTEC subsets have slightly different chromatin structures. Cluster 3 was mostly transferred to cluster R1 (88.2%) (Figure 3B and C), also designated as Aire^+^ mTECs. Interestingly, cells transferred to cluster R9 are embedded around the junction between cluster 0 and 3 (Figure 3B), suggesting that cluster R9 may be a transitional stage between R1 and R0. Clusters 1, 2, and 6 are closely embedded in the UMAP dimension and principally assigned to clusters R2, R4, and R5 (Figure 3B and C), suggesting that these clusters are concordant with mTEC I or intertypical TECs assigned in the scRNA-seq data. Cluster 4 mainly contains cells transferred to cluster R7 (55.0%) and R10 (27.2%) (Figure 3B), which were assigned as post-Aire mTECs (mTEC III). Cells assigned in R7 and R10 were embedded in distinct regions of cluster 4, implying that post Aire^+^ mTECs consist of two cell types with slightly different gene expression profiles and chromatin structures. As expected, cluster 5 with an open *Lrmp* gene was transferred to cluster R6, a tuft-like mTEC subset (mTEC IV). Cluster 9 was assigned as cluster R13, which was Gp2^+^ TECs (Figure 3B and C). Cluster 7 was transferred to cluster R8 and R12, assigned as cTECs and jTECs, respectively. Cluster 8 contains clusters R15 (64.7%) and R16 (34.2%), which express markers of structural TECs and cilia TECs, respectively (Figure 3C and Figure 3—figure supplement 1). Finally, cluster 10 was transferred to clusters R11 and R14, which are assigned as T cells and Nurse TECs (Figure 3C and Figure 3—figure supplement 1). Although a few cells were transferred to R17 in scRNA-seq, these cells did not form cluster in this analysis. Thus, TEC heterogeneity predicted from scRNA-seq may be ascribed to differences in chromatin structure.

**Figure 3.**
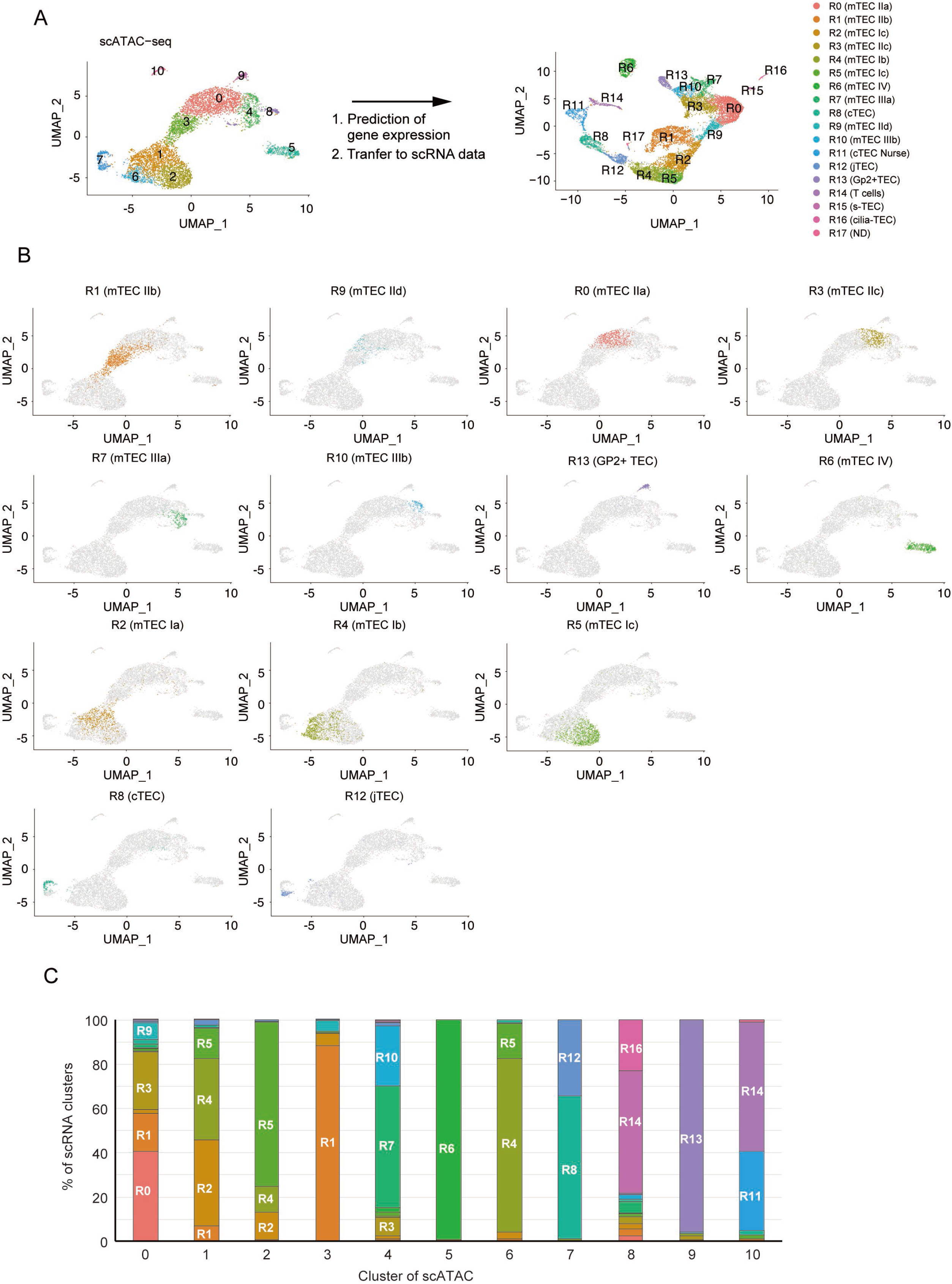
Integrative analysis of scATAC-seq data and scRNA-seq data of TECs. **A**. Gene expression was predicted from scATAC-seq data using Signac. Individual cells in the cluster from scATAC-data (cluster 0 to were assigned and transferred to the UMAP plot of scRNA-seq cluster (R0 to R17). **B**. Correlation between clusters derived from scATAC-seq and scRNA-seq datasets of TECs. Cell types were annotated in scATAC-data set of TECs by transferring clusters from an scRNA-seq dataset. **C**. Ratio of cells assigned to each scRNA-seq cluster in each scATAC cluster.

### Aire-positive mTECs are divided into two subsets having distinct chromatin structures

Previous scRNA-seq studies proposed the existence of a TEC population showing high expression of cell-cycle-regulated genes^25, 26, 31^. In our scRNA-seq data, cluster R1 (mTEC IIb) appears equivalent to such a TEC subset, expressing cell-cycle-related genes (Figure 2—figure supplement 1). Sub-clustering of cluster R1 showed its separation into 5 sub-clusters (R1A to R1E in Figure 2—figure supplement 3A and B). Clusters R1A, R1B, R1C and R1D showed expression of *Aire*. In contrast, *Ccl21a*, but not *Aire*, is highly expressed in cluster R1E (Figure 2—figure supplement 3C and D). This is largely consistent with a previous study. Thus, TECs expressing cell-cycle-related genes defined in scRNA-seq analysis may be divided into Aire-positive and Aire-negative *Ccl21a*^high^ subsets^31^.

Integrative analysis of scRNA-seq and scATAC-seq suggested that cells in cluster 3 in scATAC-seq were transferred to cluster R1. Notably, although both clusters 0 and 3 have the accessible enhancer element of the Aire gene (Figure 1), 327 genomic regions were significantly opened, and 85 regions were closed in cluster 3, in contrast to cluster 0 (Supplementary Table 2 and Figure 2—figure supplement 1B). Thus, it is likely that the Aire^+^ mTECs subset expressing cell cycle-related genes have a distinct chromatin structure relative to other TEC subsets.

Notably, some cells of clusters 1 (7%) and 0 were assigned as cluster R1 (Figure 3). This may be consistent with the heterogeneity of R1, suggested from the subcluster analysis (Figure 2—figure supplement 3). Consistently, scATAC-seq analysis showed that chromatin accessibility of a marker gene for cluster R1E (*Mgp*, Figure 2—figure supplement 3B) was low in cluster 3 and relatively higher in cluster 1 (Figure 2—figure supplement 3E). Thus, this analysis suggests that R1 includes a *Ccl21a*^high^ TEC subset having a chromatin structure different from the Aire^high^ TEC subsets in cluster R1. Thus, it is possible that TECs expressing cell-cycle-related genes, proposed by scRNA-seq analysis, contain at least two proliferating TECs subsets having different chromatin accessibilities and gene expression profiles.

RNA velocity, which recapitulates differentiation dynamics by comparing unspliced and spliced RNA in scRNA-seq data^41^, predicted that cluster R1 may differentiate into other Aire^+^ mTECs (clusters R0, R3 and R9) (Figure 3—figure supplement 2A), which is consistent with analyses of others^25^. Moreover, trajectory analysis of scATAC-seq data using the Monocle3 package also suggested a possible transition between cluster 3 and cluster 0 (Figure 3—figure supplement 2B). Thus, these trajectory analyses of scRNA-seq and scATAC-seq suggest that the Aire^+^ mTEC subset expressing cell-cycle-related genes may be precursors of other Aire^+^ mTECs. Thus, integrative analysis of scATAC-seq and scRNA-seq data imply that cluster 3 (cluster R1) may be equivalent to transiently amplifying cells (TA cells) with a distinct chromatin structure.

### A proliferative cell subset is present in Aire^+^ mTECs

TA cells were defined as a proliferative, short-lived cell subset generated from progenitor or stem cells and differentiating into mature quiescent cells^27, 42^. To search for evidence supporting the presence of TA cells of mTECs (TA-TECs), we first sought to isolate the proliferating Aire^+^ CD80^hi^ mTEC subset as candidate TA-TECs. Fucci2a mice, in which cell cycle progression can be monitored with mCherry (G1 and G0 phases) and Venus (G2, M, and S phases) fluorescence, were used to isolate such proliferating cells (Figure 4A)^32, 43, 44, 45^, and were crossed with Aire-GFP-reporter mice to facilitate detection of Aire expression^46^. Flow cytometry analysis indicated that Venus^+^ cells are present in mTECs expressing high levels of CD80 (mTEC^hi^). Moreover, these Venus^+^ mTECs^hi^ expressed Aire-GFP (Figure 4B). Thus, these data revealed the presence of dividing cells in the Aire^+^ CD80^hi^ mTEC fraction. The fluorescence intensity of Aire-GFP in Venus^+^ CD80^hi^ mTECs showed a broad peak and was slightly lower than that of Venus^-^ mTEC^hi^ cells, which may be due to the relatively lower expression of Aire in Venus^+^ CD80^hi^ mTECs. However, the compensation between GFP and Venus proteins, which have very close fluorescence spectra, hampered an exact comparison of Aire expression levels between Venus^+^ mTEC^hi^ cells and Venus^-^ mTEC^hi^ cells. We next confirmed Aire protein expression in proliferating mature mTECs. Immunostaining with an anti-Aire-antibody revealed the presence of Aire protein localized in the nucleus of sorted Venus^+^ CD80^hi^ mTECs (Figure 4C). Moreover, immunostaining of the thymic section from *Foxn1*-specific Fucci2a mice revealed that Venus^+^ cells are localized in the medulla, and some of the Aire^+^ mTECs were Venus^+^ (Figure 4D and Figure 4—figure supplement 1). Taken together, these data confirm the presence of proliferating Aire^+^ CD80^hi^ mTECs in the thymic medulla.

**Figure 4.**
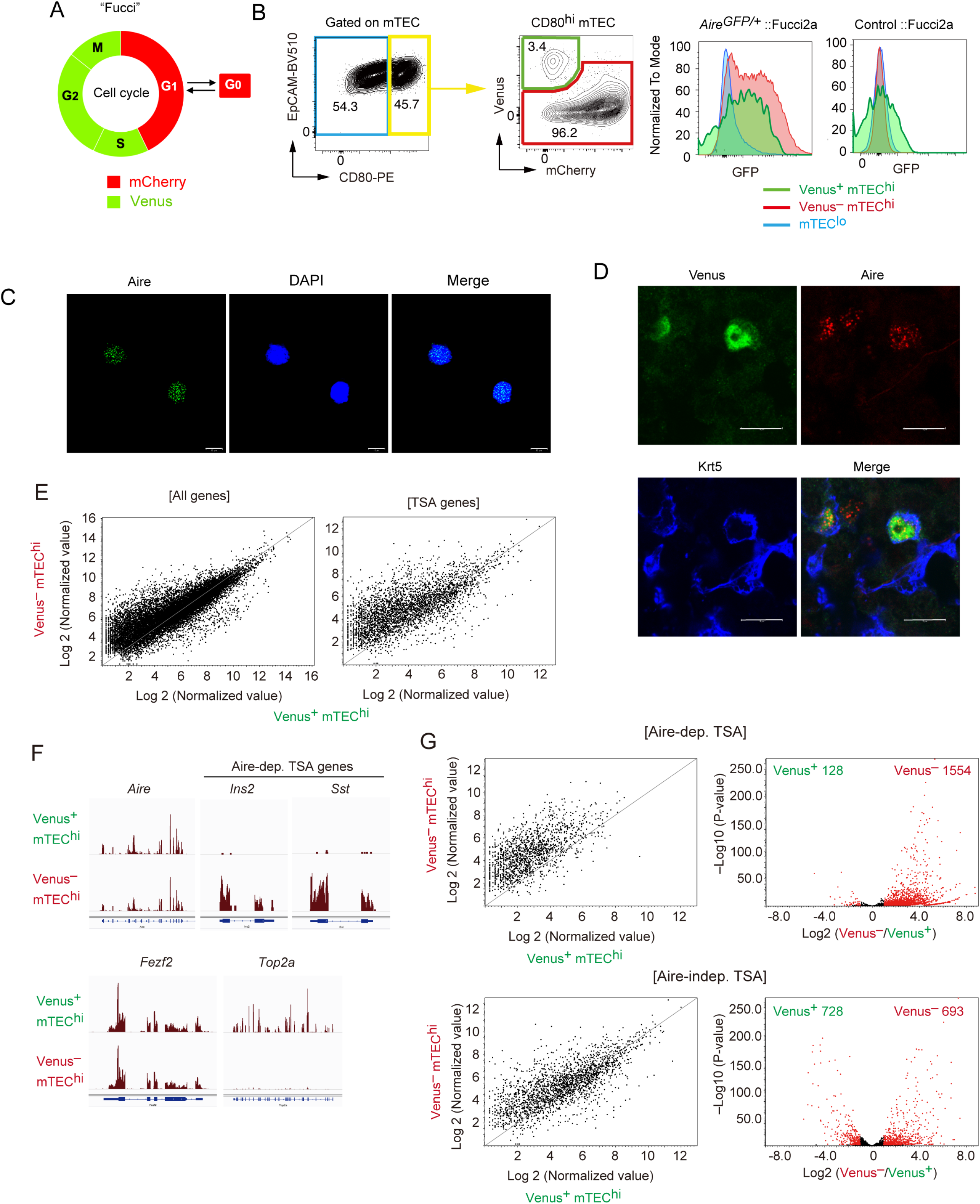
A highly proliferative subset of Aire^+^ CD80^hi^ mTECs. **A**. Schematic depiction of cell cycles and Fucci fluorescence. **B**. Flow cytometric analysis of TECs from Fucci2a mice crossed with Aire-GFP-reporter mice. The gating strategy is shown. Intensities of GFP to monitor Aire expression in each subset (Venus^+^ CD80^hi^ mTEC, Venus^-^ CD80^hi^ mTEC and CD80^lo^ mTEC^lo^) are shown in the right panels. Left, *Aire*^gfp/+^:: Fucci2a; right, control::Fucci2a. Typical figures of 3-independent experiments are exhibited. **C**. Immunostaining of a sorted Venus^+^ CD80^hi^ mTEC subset via anti-Aire antibody and DAPI (nucleus staining). Typical panels of 3-independent experiments are exhibited. Scale bars, 10 μm **D**. Immunostaining of thymic sections from Fucci2a mice with anti-Aire and anti-keratin-5 (Krt5) antibodies. Typical panels of 3-independent experiments are exhibited. Scale bars, 10 μm **E** Scatter plots of RNA sequencing data from Venus^+^ CD80^hi^ mTEC and Venus^-^ CD80^hi^ mTEC subsets. The left panel shows a plot of all detected genes and the right panel show TSA genes detected. N = 3. **F**. A typical RNA sequencing tracks of *Aire*, typical Aire-dependent TSA genes (*Ins* and *Sst*), *Fezf2*, and *Top2a* (a marker of G2/M phase). **G**. Scatter plots and volcano plots of RNA sequencing data from Venus^+^ CD80^hi^ mTEC and Venus^-^ CD80^hi^ mTEC subsets. Upper panels show Aire-dependent TSAs, lower panels show Aire-independent TSAs. Red dots in volcano plots indicate genes for which expression differed significantly (2-fold change and FDR P < 0.05) in Venus^+^ and Venus^-^ CD80^hi^ mTEC subsets. Numbers of differentially expressed genes are shown in the panels. N = 3. Y axis is log10 of FDR P-value.

### Proliferating Aire^+^ mTECs express low levels of Aire-dependent TSAs

We next addressed whether the proliferating Aire^+^CD80^hi^ mTECs subset has molecular signature distinct from that of quiescent Aire^+^CD80^hi^ mTECs. RNA-seq analysis of sorted cells from Fucci mice suggested that Venus^+^ CD80^hi^ mTECs and Venus^-^ CD80^hi^ mTECs subsets have considerably different gene expression profiles (Figure 4E). As expected, gene ontology analysis confirmed enrichment of cell cycle-related genes in Venus^+^ CD80^hi^ mTECs compared with Venus^-^ CD80^hi^ mTECs (Supplementary Table 3). Notably, although expression levels of Aire were comparable (Figure 4F), the Venus^+^ CD80^hi^ mTEC subset expressed lower levels of Aire-dependent TSAs than the Venus^-^ CD80^hi^ mTECs subset (Figure 4F and G). However, expression of Aire-independent TSAs was relatively comparable in the two subsets (Figure 4G). These data suggested that proliferating Aire^+^CD80^hi^ mTECs are phenotypically immature, compared to quiescent Aire^+^CD80^hi^ mTECs.

### Proliferating Aire^+^ mTECs are precursors of mature mTECs

Because TA cells are defined as short-lived cells differentiating into mature cells^27^, we next addressed this issue regarding the proliferating Aire^+^ CD80^hi^ mTECs. *In vivo* pulse-labeling of TECs with 5-bromo-2’-deoxyuridine (BrdU) was performed. Because mCherry^hi^ cells and mCherry^lo^ were generally in G0 and G1 stages of the cell cycle, respectively^47^, each fraction in CD80^hi^ mTECs was sorted separately after *i.p* administration of BrdU, and thereafter stained with anti-BrdU antibody (Figure 5A). This procedure was necessary because mCherry fluorescence is lost after BrdU staining. Flow cytometric analysis showed that approximately 35% of mCherry^lo^ CD80^hi^ mTECs (hereafter referred as to mCherry^lo^) were labeled at 12 h (Day 0.5) after the BrdU injection (Figure 5B). In contrast, about 3% of mCherry^hi^ CD80^hi^ mTECs (referred as to mCherry^hi^) were BrdU-positive (Figure 5B). Thus, as expected, cell cycle progression of mCherry^lo^ is much faster than mCherry^hi^. Importantly, cell number and the ratio of BrdU-positive cells in the mCherry^lo^ fraction was significantly decreased 3.5 days after the BrdU injection (Figure 5B and C). On the other hand, the frequency of BrdU-positive cells in mCherry^hi^ was increased by day 3.5, and plateaued from day 3.5 to day 6.5 (Figure 5B and C). Notably, fluorescence intensity (MFI) of BrdU staining in mCherry^hi^ at Day 3.5 was about 50% lower than that in mCherry^lo^ at Day 0.5 (Figure 5D), suggesting that mCherry^hi^ at day 3.5 were generated after cell division. Overall, these data suggest that mCherry^lo^ are transiently proliferating, and after cell division, they are converted to mCherry^hi^ having low proliferative activity.

**Figure 5.**
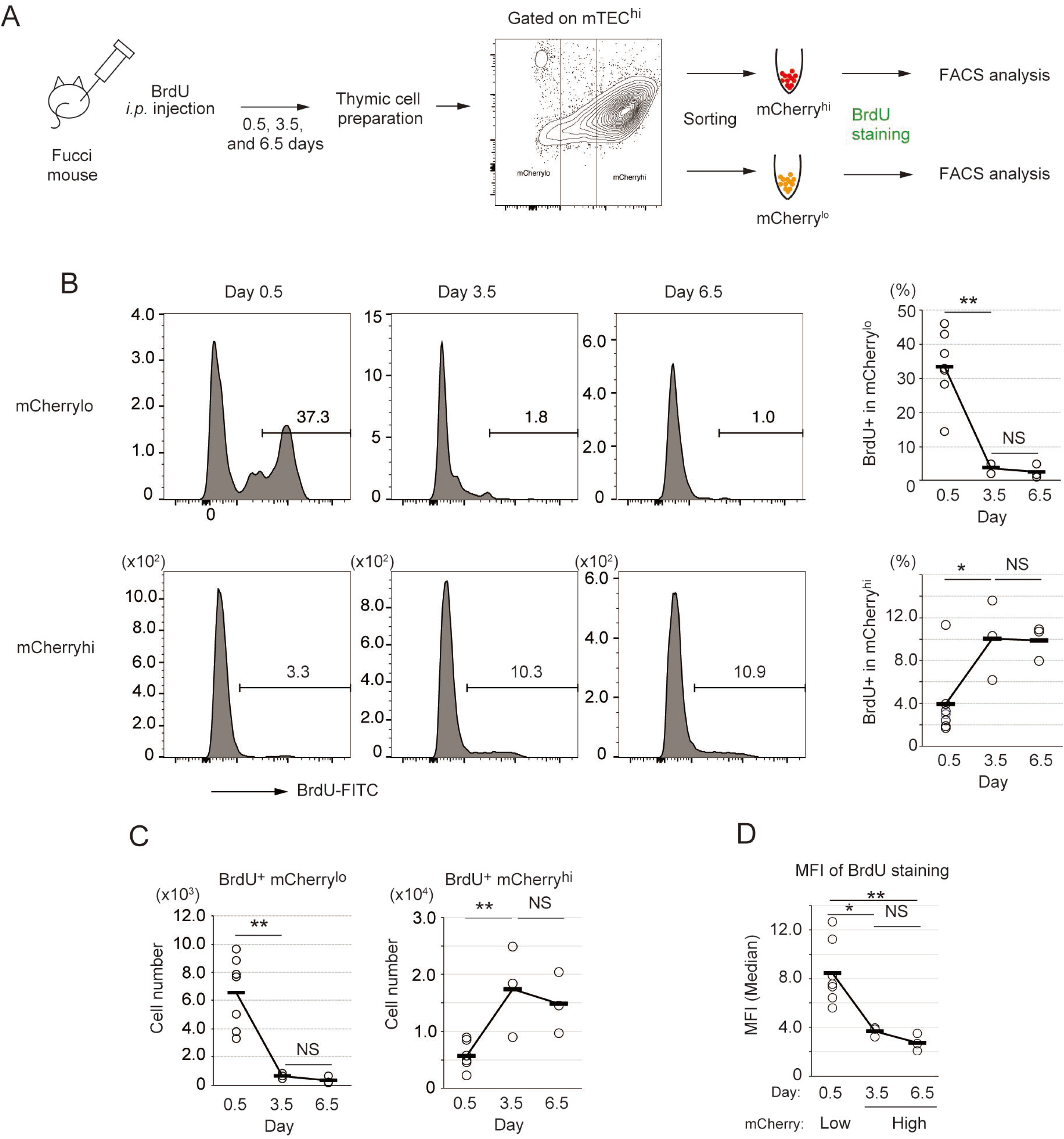
Fate mapping study by *in vivo* BrdU pulse-labeling of Fucci TECs. **A.** Schematic procedure of *in vivo* BrdU pulse-labeling of Fucci mouse, and analysis of BrdU staining in mCherry^hi^CD80^hi^ and mCherry^lo^CD80^hi^ mTECs by flow cytometiric analysis. BrdU staining was done after sorting each cell fraction. **B**. Typical flow cytometric profile of BrdU staining in mCherry^lo^CD80^hi^ mTECs (upper panels) and mCherry^hi^CD80^hi^ mTECs (lower panels) at Days 0.5, 3.5, and 6.5 after the BrdU injection. Data for the ratio of BrdU^+^ cells in each mTEC fraction are summarized in right figures. N = 7 for 0.5 day after the BrdU injection, N = 3 for 3.5 day and 6.5 day after the injection. Two-tailed Student’s t-tests. ** P < 0.01 and * P < 0.05. NS, not significant (P > 0.05). P = 1.5 x 10^-3^ for the upper figure and P = 0.033 for the lower figure. **C**. Cell number of BrdU^+^mCherry^lo^CD80^hi^ mTECs and BrdU^+^mCherry^hi^CD80^hi^ mTECs at Days 0.5, 3.5, and 6.5 after the BrdU injection. Two-tailed Student’s t-tests. ** P < 0.01. NS, not significant (P > 0.05). P = 4.3 x 10^-3^ for the left figure and P = 5.1 x 10^-3^ for the right figure. **D**. MFI of BrdU staining in mCherry^lo^CD80^hi^ at Day 0.5 and mCherry^hi^CD80^hi^ at Day 3.5 and 6.5. MFIs of other time points were difficult to evaluate because of very low cell numbers. Two-tailed Student’s t-tests. * P = 0.015 and ** P = 6.5 x 10^-3^. NS, not significant (P > 0.05).

To verify that mCherry^lo^ cells are precursors of mCherry^hi^, we performed an *in vitro* reaggregation thymic organ culture (RTOC) experiment (Figure 6A). The mCherry^lo^ fraction (Figure 6—figure supplement 1) was reaggregated with wild type embryonic thymic cells. After 5-days of culture, mCherry^hi^ was detected in RTOC (Figure 6A). Because Venus^+^mCherry^lo^ cells were practically absent in RTOC (Figure 6—figure supplement 1A), survived mCherry^lo^ cells were mostly converted into mCherry^hi^ in RTOC. Interestingly, reaggregation with allogenic fetal thymus (Balb/cA background) was not sufficient for the conversion to mCherry^hi^ (Figure 6—figure supplement 1B), implying that high affinity interaction between TCR and MHC contributes to survival and maintenance of mCherry^lo^ TECs as decribed previously^48^. Next, we sorted mCherry^hi^ cells in the RTOC (referred as to mCherry^hi^-RTOC) in addition to mCherry^lo^ and mCherry^hi^ from the Fucci thymus, and analyzed gene expression by RNA-seq. As expected, the mCherry^lo^ fraction expressed a lower level of Aire-dependent TSAs, compared to mCherry^hi^ (Figure 6B), although Aire and Mki67 were highly expressed (Figure 6C). Importantly, in comparison to the mCherry^lo^ fraction, the mCherry^hi^-RTOC fraction showed higher levels of Aire-dependent TSAs (Figure 6B). Moreover, beside cell-cycle-related genes, some genes were highly expressed in all mCherry^lo^, Venus^+^ cells, and cluster R1 cells (Figure 6—figure supplement 2 and Supplementary Table 4). Notably, these gene set were down-regulated in mCherry^hi^-RTOC (Figure 6C and Figure 6—figure supplement 2). These data suggest that mCherry^lo^ cells were converted into mCherry^hi^ in RTOC.

**Figure 6.**
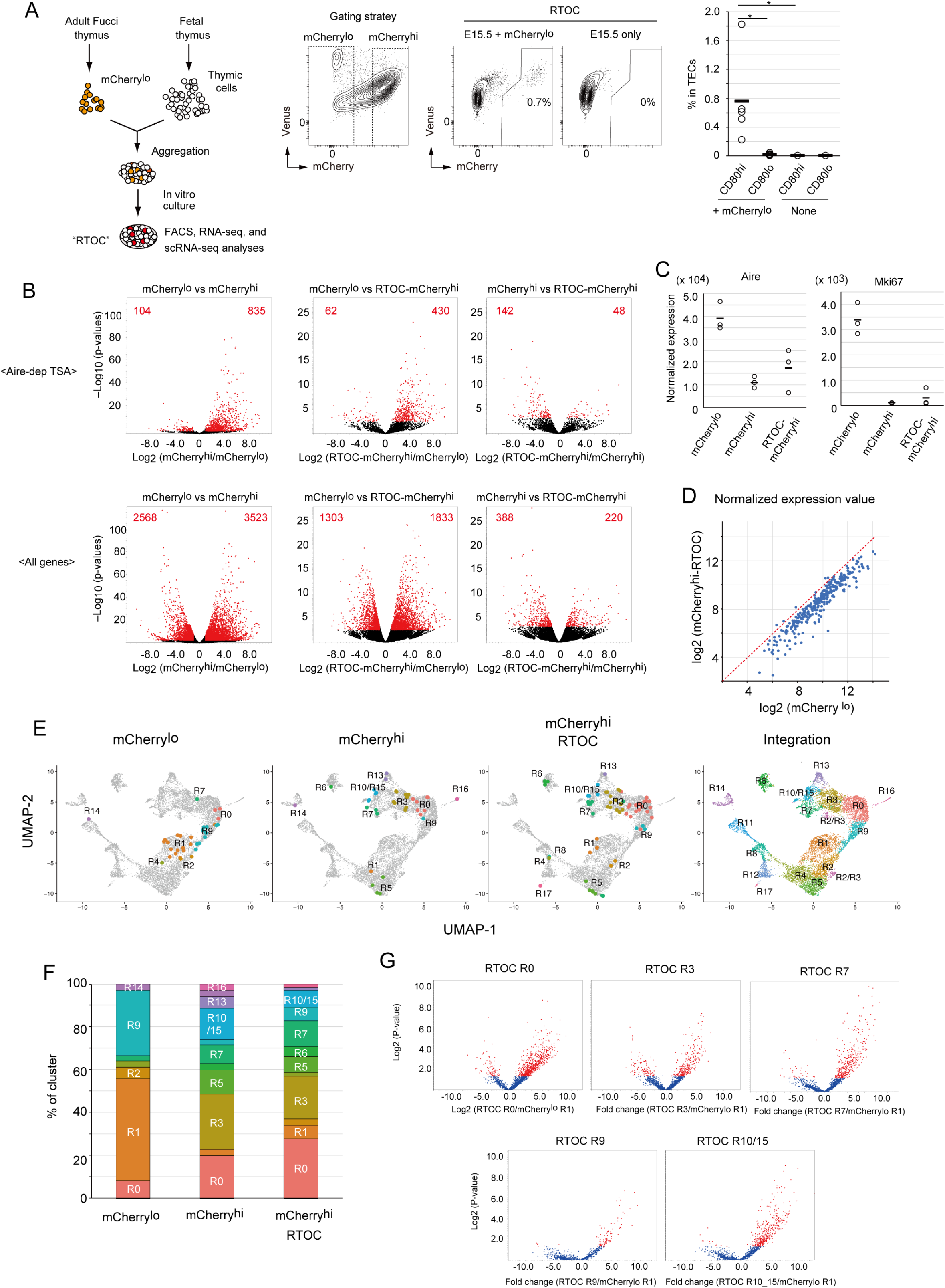
Fate mapping study of proliferating Aire^+^ mTECs in *in vitro* reaggregated thymic organ culture (RTOC) **A**. RTOC experiment to test the differentiation capacity of proliferating Aire^+^ mTECs. Proliferating Aire^+^ mTECs (mCherry^lo^) and E15.5 embryonic thymic cells were re-aggregated and subsequently cultured for 5 days. Reaggregated thymic organ (RTO) was analyzed by flow cytometer. Representative flow cytometric profiles of RTOC are shown. N = 5. The ratio of mCherry^hi^ cells in TECs is summarized in right figure. *P < 0.05. P = 0.027 between CD80^hi^ and CD80^lo^ in mCherry^lo^ and P = 0.024 between CD80^hi^ mCherry^lo^ and CD80^hi^ RTOC control. **B.** Volcano plots of RNA-seq data from mCherry^lo^ CD80^hi^ mTECs (mCherry^lo^), mCherry^hi^ CD80^hi^ mTECs (mCherry^hi^), and mCherry^hi^ CD80^hi^ mTECs in RTOC (mCherry^hi^ in RTOC). Red dots in volcano plots indicate genes for which expression differed significantly between the two subsets. Numbers of differentially expressed genes are shown in the panels. N = 3. Y axis is log10 of FDR P-value. **C.** Expression levels of Aire and Mki67 in mCherry^lo^, mCherry^hi^, and mCherry^hi^ in RTOC. **D**. Scatter plot of normalized expression values of TA-TEC marker candidates in mCherry^lo^ and mCherry^hi^ in RTOC. TA-TEC marker candidate genes were selected from bulk RNA-seq data and scRNA-seq data in Supplementary Figure 7. **E**. Integration of well-based scRamDA-seq data (mCherry^lo^, mCherry^hi^, and mCherry^hi^ in RTOC) with the droplet-based scRNA-seq data in Figure 2. **F.** Frequency of each cell cluster in scRamDA-seq data of mCherry^lo^, mCherry^hi^, and mCherry^hi^-RTOC. **G.** Volcano plot of TSA expression in each cell cluster in scRamDa-seq data of mCherry^hi^-RTOC as compared to mCherry^lo^. Red dots indicate significantly changed TSA genes.

In order to detail phenotypes of mCherry^hi^-RTOC, we next performed well-based scRNA-seq. mCherry^hi^-RTOC in addition to mCherry^lo^CD80^hi^ and mCherry^hi^CD80^hi^ mTECs from the murine thymus were single-cell sorted by flow cytometry, and then gene expression in individual cells was determined by random displacement amplification sequencing (RamDA-seq) technology^49^. After quality control of the data, gene expression matrix data of single-cell RamDA-seq (scRamDA-seq) were integrated with the droplet-based scRNA-seq data (Figure 6D). Although this integration slightly changed the UMAP dimension and clustering compared to Figure 2, assignment of each cluster was successfully achieved in the practically same fashion (Supplementary Figure 7E and F), except that cluster R15 (s-TEC) in Figure 3 was incorporated into cluster R10 (mTEC IIIb) and one new cluster were separated from cluster R2 and R3.

Cells from the mCherry^lo^CD80^hi^ mTEC fraction (total 36 cells) were assigned mainly to clusters R1 (17 cells) and R9 (11 cells) (Figure 6E and F, and Supplementary Table 5). Some cells were assigned to clusters R0 (3 cells) and R2 (2 cells). Although other cells were assigned to clusters R4, R7 and R14, the embedded position was separated from each parent cluster, which may be due to misclustering. In contrast, cells in the mCherry^hi^CD80^hi^ mTEC fraction (total 35 cells) were more heterogenous and consisted of cells assigned mainly to clusters R0 (7 cells), R3 (9 cells), R5 (4 cells), R7 (3 cells), R10/15 (5 cells), and R13 (2 cells) (Figure 6E and F, and Supplementary Table 5). Except for cluster R5, these clusters were concordant with Aire^+^ mTECs, post-Aire mTECs, and GP2^+^ TECs. Notably, after the RTOC, heterogenous cell populations including clusters R0 (18 cells), R3 (13 cells), R5 (5 cells), R6 (3 cells), R7 (8 cells) and R10/15 (5 cells) were found in the mCherry^hi^-RTOC population (total 65 cells). Its composition was relatively similar to that of the mCherry^hi^CD80^hi^ mTEC fraction (Figure 6F). Moreover, these mCherry^hi^-RTOC cells expressed high levels of TSAs (Figure 6G). Interestingly, 5 cells in mCherry^hi^-RTOC were assigned to cluster R5, which also reside in the mCherry^hi^CD80^hi^ mTEC fraction from the adult thymus. This finding is consistent with the idea of an “intertypical” mTEC cluster, which reportedly contains both CD80^hi^ mTECs and CD80^lo^mTECs^26^. Overall, these data suggest that mCherry^lo^CD80^hi^ mTECs differentiate into quiescent mature mTECs expressing high levels of TSAs, including Aire^+^ mTECs (mTEC II), post-Aire mTECs (mTEC III), and tuft-like mTECs (mTEC IV).

### Proliferating Aire^+^ mTECs are present after puberty in mice

We investigated whether proliferative Aire^+^ mTECs persisted in the thymi of older mice. TECs were analyzed in 4-week-old, 8-week-old, and 19-week-old Fucci Aire-GFP mice. Flow cytometric analysis showed that a Venus^+^ mTEC^hi^ subset was present in 19-week-old mice as well as younger mice (Figure 7A). Moreover, Venus^+^ mTEC^hi^ cells expressed Aire genes (Figure 7A). These data strongly suggested that transit amplifying TECs persist in the adult thymus as a source of mature mTECs.

**Figure 7.**
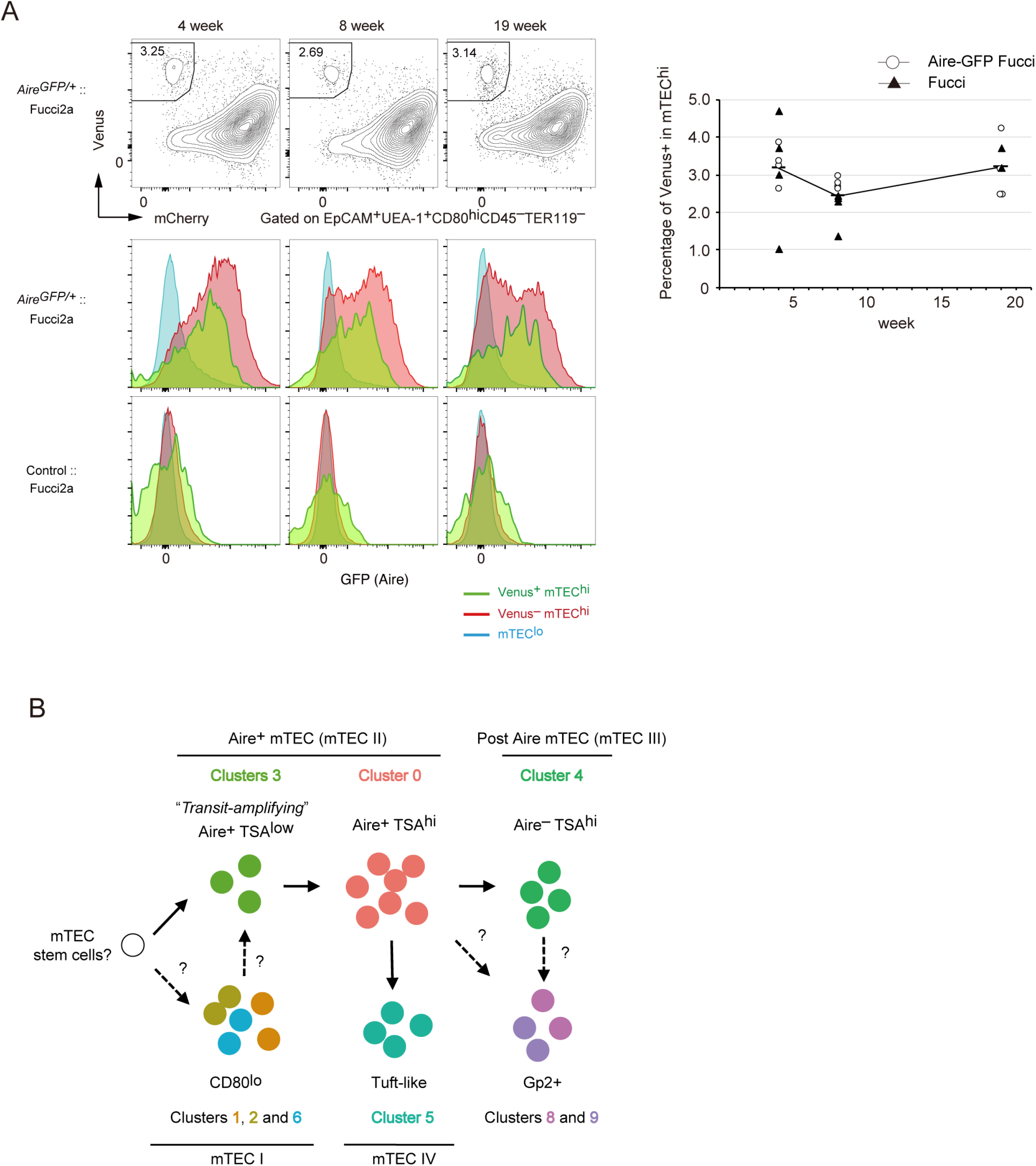
Proliferating Aire^+^ CD80^hi^ mTECs persist in older mice. **A**. Flow cytometry analysis of CD80^hi^ mTEC subsets from Fucci2a mice aged 4, 8, and 19 weeks. Representative data are shown. Percentages of Venus^+^ cells in CD80^hi^ mTEC subsets are summarized in the graph in the right panel. N = 4 each for *Aire*^gfp/+^:: Fucci2a (circle) and control::Fucci2a (closed triangles) **B**. Schematic depiction of the proposed process of Aire^+^ mTEC development in the adult thymus. Transit-amplifying TSA^lo^ Aire^+^ TECs give rise to mature mTECs. Precursor cells to the transit-amplifying TECs have not been determined yet. Cluster numbers in Figure 1 are shown together with the model of mTEC subsets I to IV.

Integrative computational analysis of our scRNA-seq data with a previously reported dataset of fetal TECs (E12 to E18) showed considerably different cell embedding between adult TECs and fetal TECs (Figure 7—figure supplement 1). A TEC-expressing subset was present in the fetal thymus whereas Aire expression was low (Clusters F3 and F12, Figure 6—figure supplement 2). This implies that fetal proliferating mTECs may have a different gene expression profile than adult proliferating Aire^+^CD80^hi^ mTECs (Figure 6—figure supplement 2).

## Discussion

With regard to mTEC differentiation in the adult thymus (Figure 7B), we hypothesize that Aire^+^ TA-TECs were generated from their Aire-negative progenitors. Aire^+^ TA-TECs (cluster 3) undergo cell division and then differentiate into quiescent Aire^+^ mTECs (cluster 0) through a transition stage, which corresponds to cluster R9 in scRNA-seq data. This differentiation process is accompanied by a chromatin structure change. Post-mitotic Aire^+^ mTECs begin high-level TSA expression, and further differentiate into post-Aire mTECs (R7, R10 and R13) by closing the Aire-enhancer region. Differentiation of mTECs expressing TSAs may have to coordinate differentiation with cell cycle regulation, as proposed in neural cells and muscle differentiation^50^.

Generally, in other tissues, transit-amplifying cells constitute a link between stem cells and mature cells^42^. An important question is what cells differentiate into proliferating Aire^+^ mTECs. Previous studies have suggested that niTECs^lo^ expressing low levels of maturation markers (i.e. CD80 or MHC II) are precursors^1, 13^. However, several recent studies have suggested that mTEC^lo^ contains several subsets, including CCL21a-positive mTECs, tuft-like mTECs, and others. One possible explanation for this is that a small number of mTEC stem cells or other precursor cells may be present in the mTEC^lo^ subset^13^. Consistently RNA velocity analysis also suggested that most mTEC^lo^ cells do not appear to differentiate into Aire-expressing mTECs. Given that transit-amplifying mTECs are present, a small number of stem/precursor cells would theoretically be sufficient for mTEC reconstitution. A previous study proposed that TECs expressing claudin 3/4 and SSEA-1 had characteristic features of mTEC stem cells in embryonic thymus^20^. We failed to detect a corresponding cluster of mTEC stem cells as a subset of adult scRNA clusters. This may be because corresponding mTEC stem cells in adult thymus are included in the “intertypical” TEC cluster, which may be a mixture of various TECs^26^. More detailed characterization of mTEC stem cells in the adult thymus is necessary to illuminate the differentiation dynamics of mTECs.

Overall, the scRNA-seq analysis in the present study suggested the presence of a novel differentiation process of TECs in the adult thymus. Disturbance of adult TEC homeostasis may cause thymoma, autoimmunity, and other diseases. Further characterization of molecular mechanisms underlying differentiation and maintenance processes in TECs will aid the development of novel therapeutic strategies against these thymus-related diseases.

## Materials and Methods

### Mice

C57BL/6 mice were purchased from Clea Japan. Littermates or age-matched, wild-type mice from the same colonies as the mutant mice were used as controls. *Aire-GFP* mice (CDB0479K, http://www2.brc.riken.jp/lab/animal/detail.php?brc_no=03515) and B6;129-Gt(ROSA)26Sor<tm1(Fucci2aR)Jkn> (RBRC06511) (Fucci2a)^32^ were provided by the RIKEN BRC through the National Bio-Resource Project of the MEXT, in Japan. CAG-Cre transgenic mice were kindly provided by Dr. Jun-ichi Miyazaki^51^. B6(Cg)-Foxn1tm3(cre)Nrm/J are from Jackson Laboratory^52^. Fucci2a mice were crossed with CAG-Cre or Foxn1-Cre mice to activate mCherry and Venus expression. Fucci mice crossed with CAG-Cre were used for all experiments except for immunostaining experiments (Figure 4). All mice were maintained under specific pathogen-free conditions and handled in accordance with Guidelines of the Institutional Animal Care and Use Committee of RIKEN, Yokohama Branch (2018-075). Almost all of available mutant and control mice were randomly used for experiments without any selection.

### Preparation of TEC suspensions and flow cytometry analysis

Murine thymi were minced using razor blades. Thymic fragments were then pipetted up and down to remove lymphocytes. Then fragments were digested in RPMI 1640 medium containing Liberase™ (Roche, 0.05 U/mL) plus DNase I (Sigma-Aldrich) via incubation at 37°C for 12 min three times. Single-cell suspensions were stained with anti-mouse antibodies. Dead cells were excluded via 7-aminoactinomycin D staining. Cells were sorted using a FACS Aria instrument (BD). Post-sorted cell subsets were determined to contain > 95% of the relevant cell types. Data were analyzed using Flowjo 10. No data points or mice were excluded from the study. Randomization and blinding were not used.

### Droplet-based scRNA-seq analysis

For scRNA-seq analysis, cell suspensions of thymi from 3 mice were prepared and pooled for each individual scRNA-seq experiment. Two experiments were performed. Cellular suspensions were loaded onto a Chromium instrument (10× Genomics) to generate a single-cell emulsion. scRNA-seq libraries were prepared using Chromium Single Cell 3’ Reagent Kits v2 Chemistry and sequenced in multiplex on the Illumina HiSeq2000 platform (rapid mode). FASTQ files were processed using Fastp^53^. Reads were demultiplexed and mapped to the mm10 reference genome using Cell Ranger (v3.0.0). Processing of data with the Cell Ranger pipeline was performed using the HOKUSAI supercomputer at RIKEN and the NIG supercomputer at ROIS National Institute of Genetics. Expression count matrices were prepared by counting unique molecule identifiers. Downstream single-cell analyses (integration of two datasets, correction of dataset-specific batch effects, UMAP dimension reduction, cell cluster identification, conserved marker identification, and regressing out cell cycle genes) were performed using Seurat (v3.0)^54^ Briefly, cells that contained a percentage of mitochondrial transcripts > 15% were filtered out. Genes that were expressed in more than 5 cells and cells expressing at least 200 genes were selected for analysis. Two scRNA-seq datasets were integrated with a combination of Find Integration Anchors and Integrate Data functions^55^. To investigate the effects of regressing out cell cycle genes on cell clustering, we compared three types of pre-processing; no regressing out, regressing out the difference between the G2/M and S phase scores, and complete regressing out of all cell cycle scores (Supplementary Fig. S3) after assigning cell cycle scores via the Cell Cycle Scoring function. The murine cell cycle genes equivalent to human cell cycle genes listed in Seurat were used for assigning cell cycle scores.

For comparison with a previously reported RNA sequence dataset obtained via a well-based study^23^, the expression matrix of unique molecule identifiers was used. Integration of the two datasets was performed using the Seurat package as described above. RNA velocity analysis was performed using velocyto. Bam/sam files obtained from the Cell Ranger pipeline were transformed to loom format on velocyto.py. RNA velocity was estimated and visualized using loom files by the velocyto R package and pagoda2.

### Droplet-based scATAC-seq analysis

In scRNA-seq analysis cell suspensions of thymi from 3 mice were prepared and pooled for each individual scRNA-seq experiment. EpCAM^+^CD45^-^TER119^-^ fraction was collected by using cell sorter (BD Aria). After washing with PBS containing 0.04% BSA, sorted cells were suspended in lysis buffer containing 10mM Tris-HCl (pH 7.4), 10 mM NaCl, 3 mM MgCl_2_, 0.1% Tween-20, 0.1% NP-40, 0.01% Digitonin, and 1% BSA on ice for 3 min. Wash buffer containing 10mM Tris-HCl (pH 7.4), 10 mM NaCl, 3 mM MgCl_2_, 0.1% Tween-20, and 1% BSA was added to the lysed cells. After centrifuging the solution, a nuclear pellet was obtained by removing the supernatant and the pellet was re-suspended in wash buffer. The concentration of nuclei and their viability were determined by staining with acridine orange/propidium iodide, and 10,000 nuclear suspensions were loaded onto a Chromium instrument (10× Genomics) to generate a single-cell emulsion. scATAC-seq libraries were prepared using Chromium Next GEM Single Cell ATAC Reagent Kits v1.1 and sequenced in multiplex on an Illumina Hiseq X ten platform. Reads were demultiplexed and mapped to the mm10 reference genome with Cell Ranger ATAC. Processing data with the Cell Ranger pipeline was performed using the NIG supercomputer at ROIS National Institute of Genetics. Downstream single-cell analyses (integration of two datasets, correction of dataset-specific batch effects, UMAP dimension reduction, cell cluster identification, conserved marker identification, and regressing out cell cycle genes) were performed using Seurat (v3.0)^54^. Briefly, cells that contained a percentage of mitochondrial transcripts > 15% were filtered out. Genes that were expressed in more than 5 cells and cells expressing at least 200 genes were selected for analysis. Two scRNA-seq datasets were integrated with a combination of Find Integration Anchors and Integrate Data functions^55^. To investigate the effects of regressing out cell cycle genes on cell clustering, we compared three types of pre-processing; no regressing out, regressing out the difference between the G2/M and S phase scores, and complete regressing out of all cell cycle scores after assigning cell cycle scores via the Cell Cycle Scoring function. The murine cell cycle genes equivalent to human cell cycle genes listed in Seurat were used for assigning cell cycle scores.

### Well-based scRNA-seq analysis

Single-cells were sorted into PCR tubes containing 1μl of cell lysis solution (1:10 Cell Lysis buffer(Roche), 10U/μl Rnasin plus Ribonuclease inhibitor (Promega) using a cell sorter, shaken at 1400 rpm for 1 min with a thermo mixer, and then stored at −80°C. Cell lysates were denatured at 70 °C for 90 s and held at 4 °C until the next step. To eliminate genomic DNA contamination, 1 μL of genomic DNA digestion mix (0.5× PrimeScript Buffer, 0.2 U of DNase I Amplification Grade, in RNase-free water) was added to 1μL of the denatured sample. The mixtures were mixed by gentle tapping, incubated in a T100 thermal cycler at 30 °C for 5 min and held at 4 °C until the next step. One microliter of RT-RamDA mix (2.5×PrimeScript Buffer, 0.6 pmol oligo(dT)18, 8 pmol 1st-NSRs, 100 ng of T4 gene 32 protein, and 3× PrimeScript enzyme mix in RNase-free water) was added to 2 μL of the digested lysates. The mixtures were mixed with gentle tapping, and incubated at 25 °C for 10 min, 30 °C for 10 min, 37 °C for 30min, 50 °C for 5 min, and 94 °C for 5 min. Then, the mixtures were held at 4 °C until the next step. After RT, the samples were added to 2 μL of second-strand synthesis mix containing 2.25× NEB buffer 2 (NEB), 0.625mM each dNTP Mixture (NEB), 40 pmol 2nd-NSRs, and 0.75 U of Klenow Fragment (NEB) in RNase-free water. Mixtures were again mixed by gentle tapping, and incubated at 16°C for 60 min, 70°C 10 min and then at 4 °C until the next step. The above-described double-stranded cDNA was purified using 15 μl of AMPure XP SPRI beads (Beckman Coulter) diluted 2-fold with Pooling buffer (20% PEG8000, 2.5 M NaCl, 10 mM Tris-HCl pH8.0, 1 mM EDTA, 0.01% NP40) and Magna Stand (Nippon Genetics). Washed AMPure XP beads attached to double-stranded cDNAs were directly eluted using 3.75 μL of 1x Tagment DNA Buffer (10 mM Tris-HCl pH8.5, 5 mM MgCl_2_, 10% DMF) and mixed well using a vortex mixer and pipetting. Diluted Tn5-linker complex was added to the eluate and the mixture was incubated at 55°C for 10 min, then 1.25μl of 0.2% SDS was added and incubated at room temperature for 5 min. After PCR for adoptor ligation, sequencing library DNA was purified using 1.0× the volume of AMPure XP beads and eluted into 24 μL of 10 mM Tris-Cl, pH 8.5.

### Standard RNA sequencing analysis

Total RNA was prepared using TRIzol reagent (Thermo Fisher Scientific) in accordance with the manufacturer’s protocol. After rRNA was depleted using the NEBNext rRNA Depletion Kit, the RNA sequence library was prepared using the NEBNext Ultra Directional RNA Library Prep Kit (New England Biolabs). Paired-end sequencing was performed with NextSeq500 (Illumina). Sequence reads were quantified for annotated genes using CLC Genomics Workbench (Version 7.5.1; Qiagen). Gene expression values were cut off at a normalization expression threshold value of 3. Differential expression was assessed via empirical analysis with the DGE (edgeR test) tool in CLC Main Workbench, in which the Exact Test of Robinson and Smyth was used^56^. An FDR-corrected *p* value was used for testing statistics for RNA-sequencing analysis. Previously described lists of TSAs and Aire-dependent TSAs^21^ were used for the analysis.

### RTOC and RNA-seq analysis

mCherry^lo^ cells (4 x 10^4^ ~ 1 x 10^5^) were sorted from Fucci mice and subsequently re-aggregated with trypsin-digested thymic cells (1 ~ 2 x 10^6^) from E15.5 wild-type mice. RTOCs were cultured on Nucleopore filters (Whatman) placed in R10 medium containing RPMI1640 (Wako) supplemented with 10% fetal bovine serum (FBS), 2 mM L-glutamine (Wako), 1× nonessential amino acids (NEAAs; Sigma-Aldrich), 0.1 pM cholera Toxin Solution (Wako 030-20621), 5 μg/ml Insulin solution from bovine pancreas (SIGMA I0516-5ML), 2 nM Triiodo-L-thyronine (SIGMA T2877-100MG),1000 units/ml LIF (nacalai NU0012-1), 0.4 μg/ml hydrocortisone,10 ng/ml EGF (Gibco PMG8041), 1 μg/ml RANKL (Wako), penicillin-streptmycin mixed solution (Nacalai Tesque), and 50 μM 2-mercaptoethanol (Nacalai Tesque) for 5 days. For RNA-seq of RTOC experiments, random displacement amplification sequencing (RamDA-seq) were used^49^, which allows RNA-seq analysis of low numbers of cells. Briefly, sorted cells were lysed in TCL buffer (Qiagen). After purification of nucleic acids by Agencourt RNA Clean XP (Beckman Coulter) and subsequent treatment with DNase I, the RT-RamDA mixture containing 2.5x PrimeScript Buffer (TAKARA), 0.6 μM oligo(dT)18 (Thermo), 10 μM 1^st^ NSR primer mix, 100 μg/mL of T4 gene 32 protein, and 3× PrimeScript enzyme mix (TAKARA) were added to the purified nucleic acids for reverse transcription. Samples were added to second-strand synthesis mix containing 2× NEB buffer 2 (NEB), 625 nM dNTP Mixture (NEB), 25 μM 2^nd^ NSR primers, and 375 U/mL of Klenow Fragment (3’-5’ exo-) (NEB). After cDNA synthesis and subsequent purification by AMPure XP (Beckman Coulter), sequencing library DNA was prepared using the Tn5 tagmentation-based method. Single-read sequencing was performed using a HiSeq2500 (v4, high out mode). Sequence reads were quantified for annotated genes using CLC Genomics Workbench (Version 7.5.1; Qiagen).

### Immunohistochemistry

The thymus was fixed with 4% paraformaldehyde and frozen in OCT compound. After washing cryosections (5 μm) with PBS, sections were blocked with 10% normal goat serum. Keratin-5 was detected using a combination of a polyclonal rabbit anti-mouse keratin-5 antibody (1:500) and AlexaFluor647-donkey-anti-rabbit IgG. Aire was detected using a labeled monoclonal antibody (1:300). Confocal color images were obtained using a LAS X (Leica) microscope.

### Immunocytochemistry

Thymic cell suspensions prepared via Liberase™ digestion were stained with anti-CD45-PE and anti-TER119-PE. After depletion of labeled CD45^+^ and TER119^+^ cells via anti-PE microbeads and a magnetic-activated cell sorting separator, negatively selected cells were stained with anti-EpCAM (CD326), anti-CD80, anti-Ly51, and UEA-1. Venus^+^ CD80^hi^ mTECs were sorted and spun down on glass slides using a Cytospin. The slides were then fixed with acetone and stained with anti-Aire antibody and DAPI for nuclear staining. Confocal images were obtained using an LAS X microscope.

### Statistical analysis

Statistically significant differences between mean values were determined using Student’s t-test (***P < 0.001, **P < 0.01 and * P < 0.05). Principle component analysis was performed using the prcomp function in R-project. The sample size was not predetermined by statistical methods but based on common practice and previous studies^6, 57^. All replicates are biological replication. All outliers were included in data.

## Data availability

FASTQ data of RNA-Seq and ATAC-seq are deposited in DDBJ (DRA009125 DRA010209 DRA12308, DRA12309 and DRA012452). Any additional information required to reanalyze the data reported in this paper was up-loaded as a zip file of Source_data.

## Acknowledgments

This work was supported by Grants-in-Aid for Scientific Research from JSPS (17H04038, 17K08622, 20K07332, 20H03441) (T.A., N.A.), grants from the Princess Takamatsu Cancer Research Fund (T.A.), The Uehara Memorial Foundation (T.A.), and The Novartis Foundation for the Promotion of Science (T.A.), and a Grant-in-Aid for Scientific Research in Innovative Areas from MEXT (18H04989, 19H04821) (T.A., N.A.). CREST from Japan Science and Technology Agency (JPMJCR2011) (T.A.). The authors declare no competing financial interests. We thank the sequencing staff at the RIKEN Center for Integrative Medical Sciences for assisting with RNA-seq. Computations were partially performed on the NIG supercomputer at ROIS, National Institute of Genetics.

## Author contributions

T.M, MM, TI, MY, HY, AI, and EO performed experiments and analyzed data. TK, TWT, SH, KH, YT, and TS analyzed data. HI and NY established mutant mouse lines. ASS, AM, and AK contributed to data analysis and interpretation. NA and TA designed the study, analyzed data, and wrote the manuscript.

---

**Supplementary Table 1**. Integration of scRNA-seq cluster and scATAC-seq cluster

**Supplementary Table 2**. Open regions in cluster 3 as compared to cluster 0 in scATAC-seq.

**Supplementary Table 3**. GO analysis of genes differentially expressed in Venus+ cells

**Supplementary Table 4.** Possible marker gene candidates for transit amplifying TECs.

**Supplementary Table 5**. Summary for assignment of individual single cells in scRamDa-seq of mCherry^hi^, mCherry^lo^, and mCherry^hi^-RTOC

**Figure 1-figure supplement 1.**
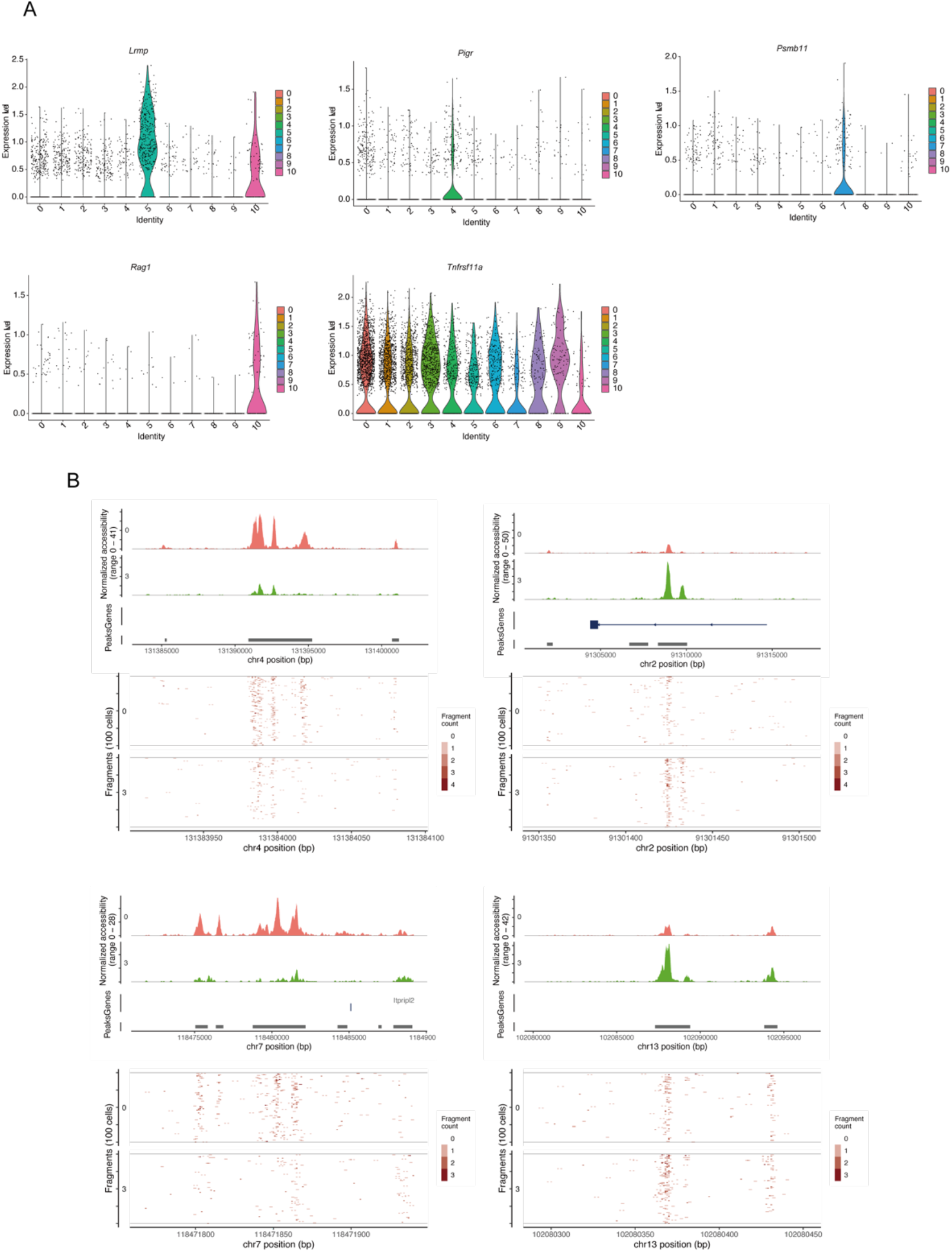
**A**. Violin plot of chromatin accessibility in TEC marker gene regions in each cluster. **B**. Pseudo-bulk accessibility tracks and frequency of sequenced fragments. Typical differentially accessible regions between clusters 0 and 3 are depicted from Supplementary Table 1.

**Figure 2-figure supplement 1.**
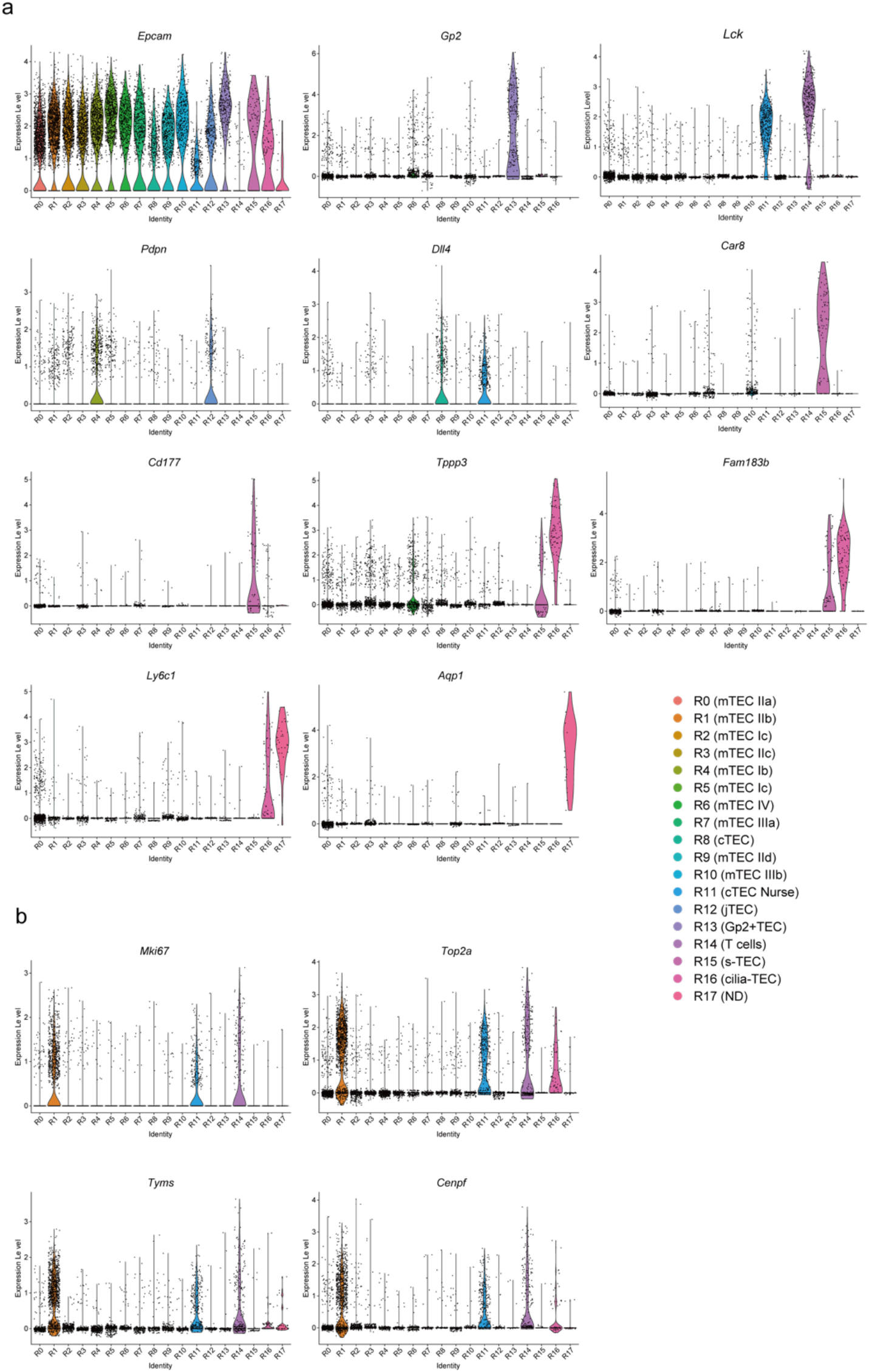
**A**. Violin plots for expression level of typical TEC marker genes in scRNA-seq analysis of TECs **B**. Violin plots for expression level of cell-cycle-related genes in scRNA-seq analysis of TECs

**Figure 2-figure supplement 2.**
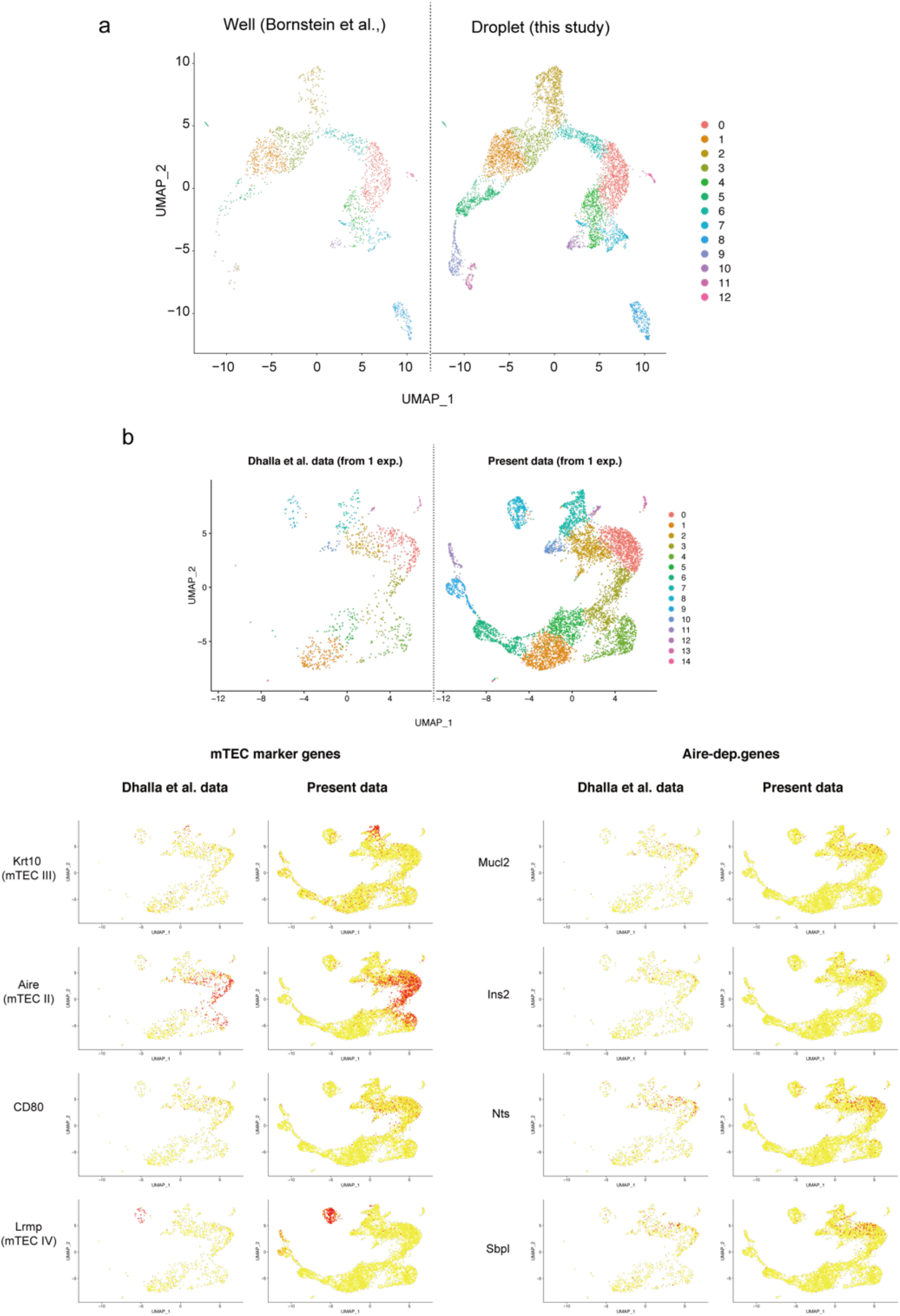
**A**. Integration of scRNA-seq data derived from a previously reported well-based study^1^ and scRNA-seq data derived from the present droplet-based study. scRNA-seq data from the two studies were integrated **B.** UMAP projections of the two scRNA-seq datasets are shown. Data from a previous study^2^ were re-analyzed and integrated with data from the present study. Expression of typical marker genes in each data.

**Figure 2-figure supplement 3.**
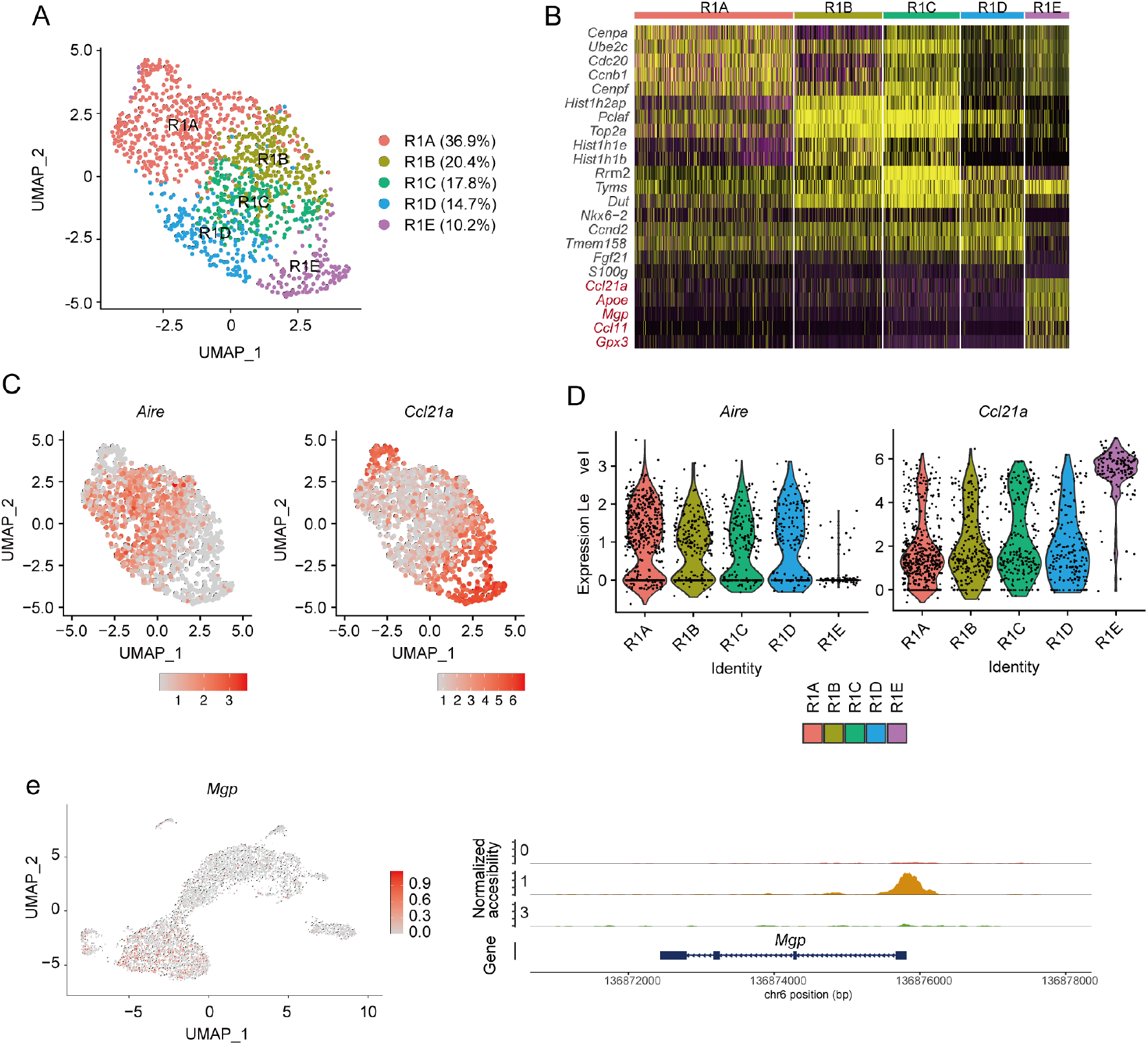
Sub-cluster analysis of the TEC subset expressing a high level of cell-cycle-related genes. **A.** UMAP plot of scRNA-seq data and the percentage of each cluster (R1A to R1E) in R1. **B**. Heatmap of the top 5 genes selectively expressed in each cluster. Yellow color indicates high expression. **C**. Expression levels of *Aire* and *Ccl21a* a in the sub-cluster is shown in dot plots. **D**. Expression levels of *Aire* and *Ccl21a* in the sub-cluster is exhibited as violin plots. **E**. Chromatin accessibility of a typical marker gene for sub-cluster RIE (*Mgp*). Accessibility in *Mgp* gene regions is represented in red (left). Pseudo-bulk accessibility tracks for *Mgp* in cluster 0, 1 and 3 is exhibited (bottom)

**Figure 3-figure supplement 1.**
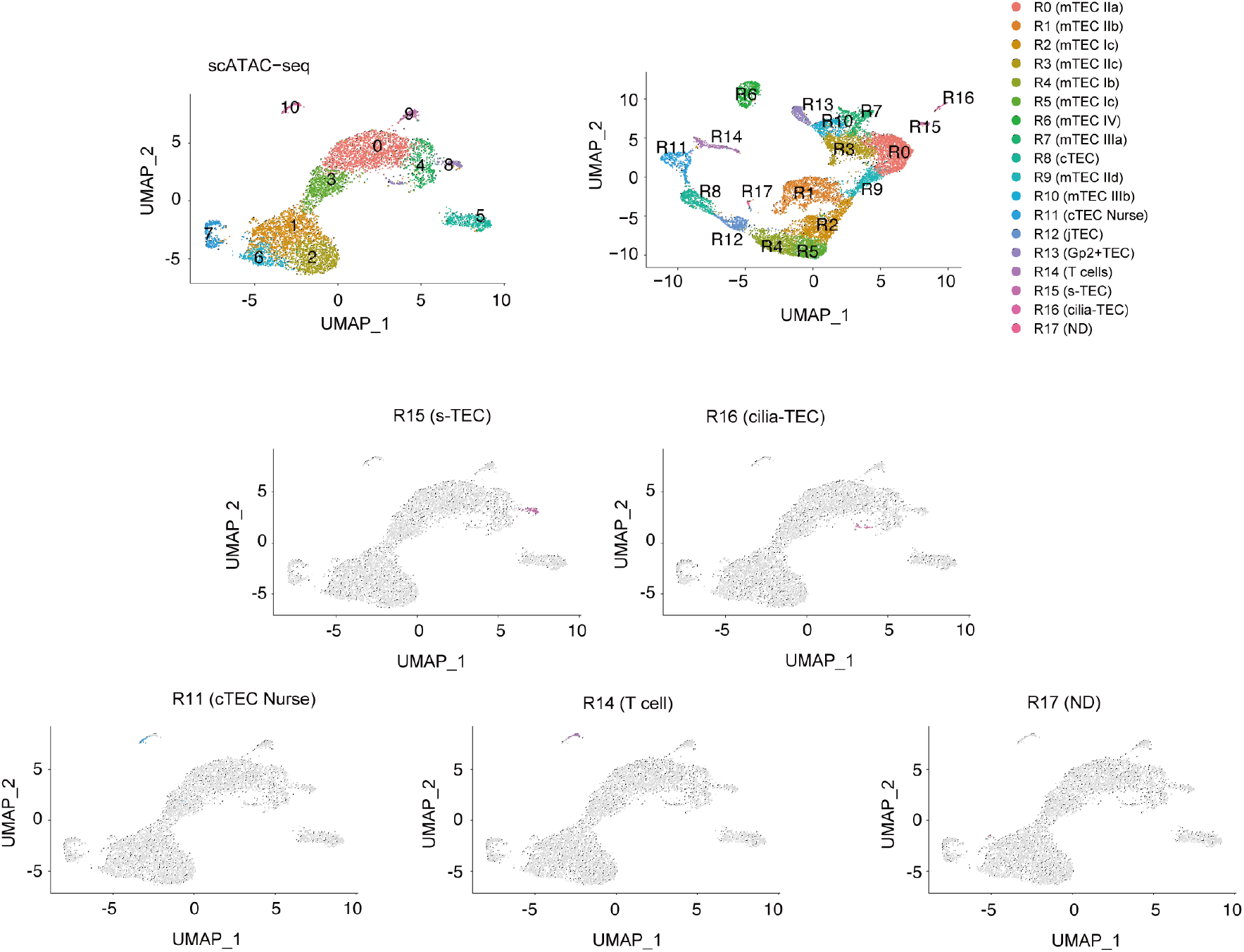
Integrative analysis of scATAC-seq data and scRNA-seq data of TECs. Gene expression was predicted from individual cells in scATAC-seq data (clusters 9 and 10). Individual cells in the scATAC-data were assigned to a scRNA-seq cluster (R0 to R17).

**Figure 3-figure supplement 2.**
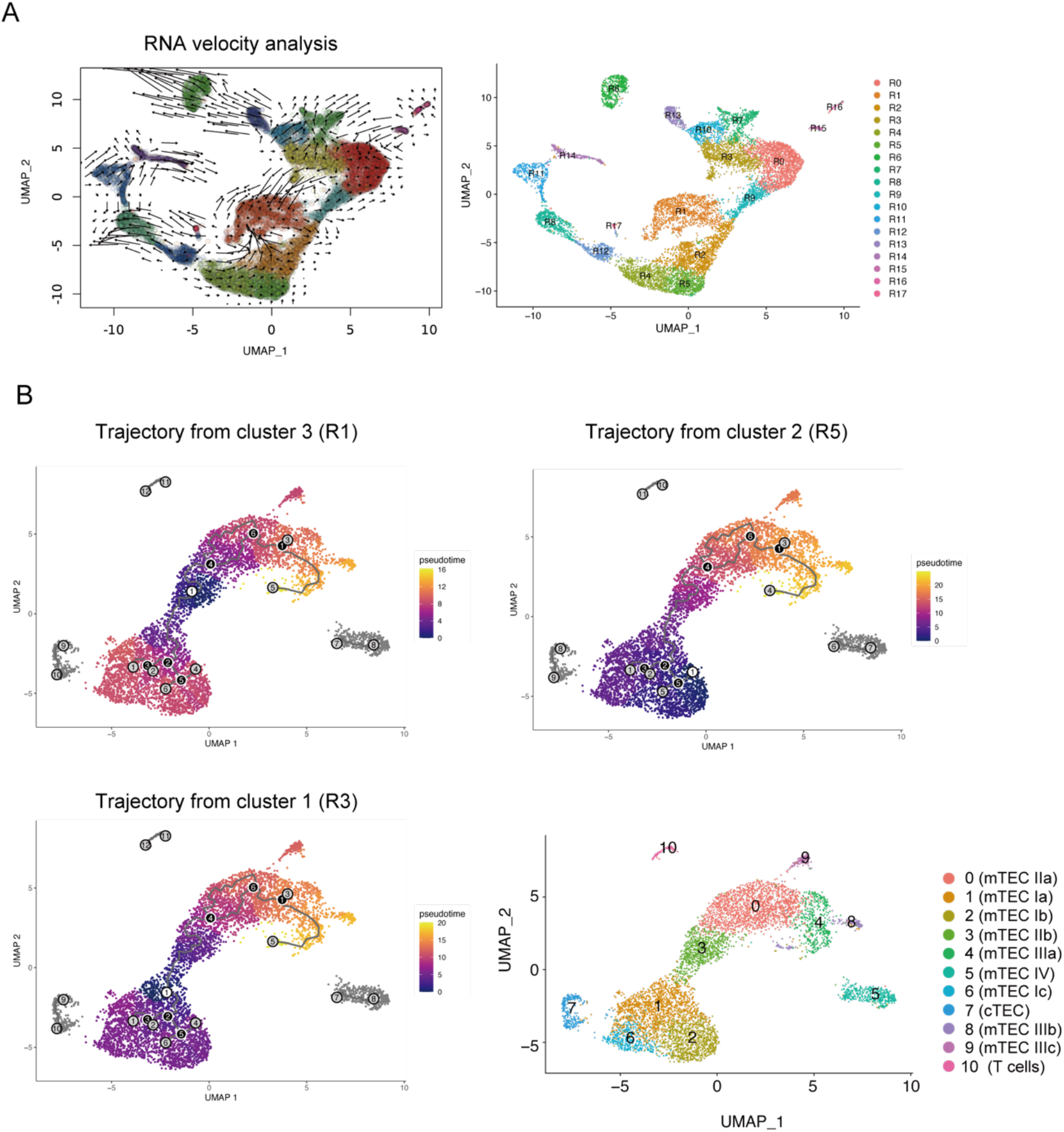
**A**. RNA velocity analysis of scRNA-seq data. **B**. Monocle 3 trajectory analysis of scATAC-seq data. The trajectory was manually started from cluster 3, 2 or 1.

**Figure 4-figure supplement 1.**
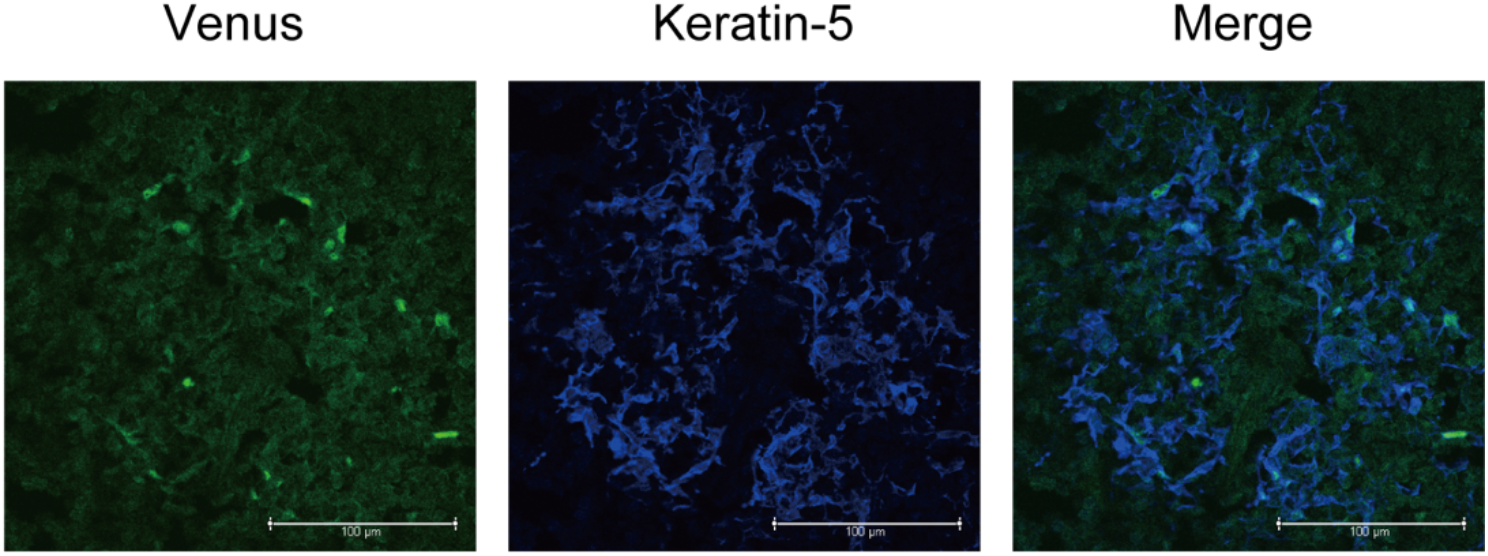
Immunostaining of thymic sections from Fucci2a mice with anti-GFP (for Venus staining, green) and anti-keratin-5 (Krt5, blue) antibodies. Typical panels of 3-independent experiments are exhibited. Scale bars, 100 μm.

**Figure 6-figure supplement 1.**
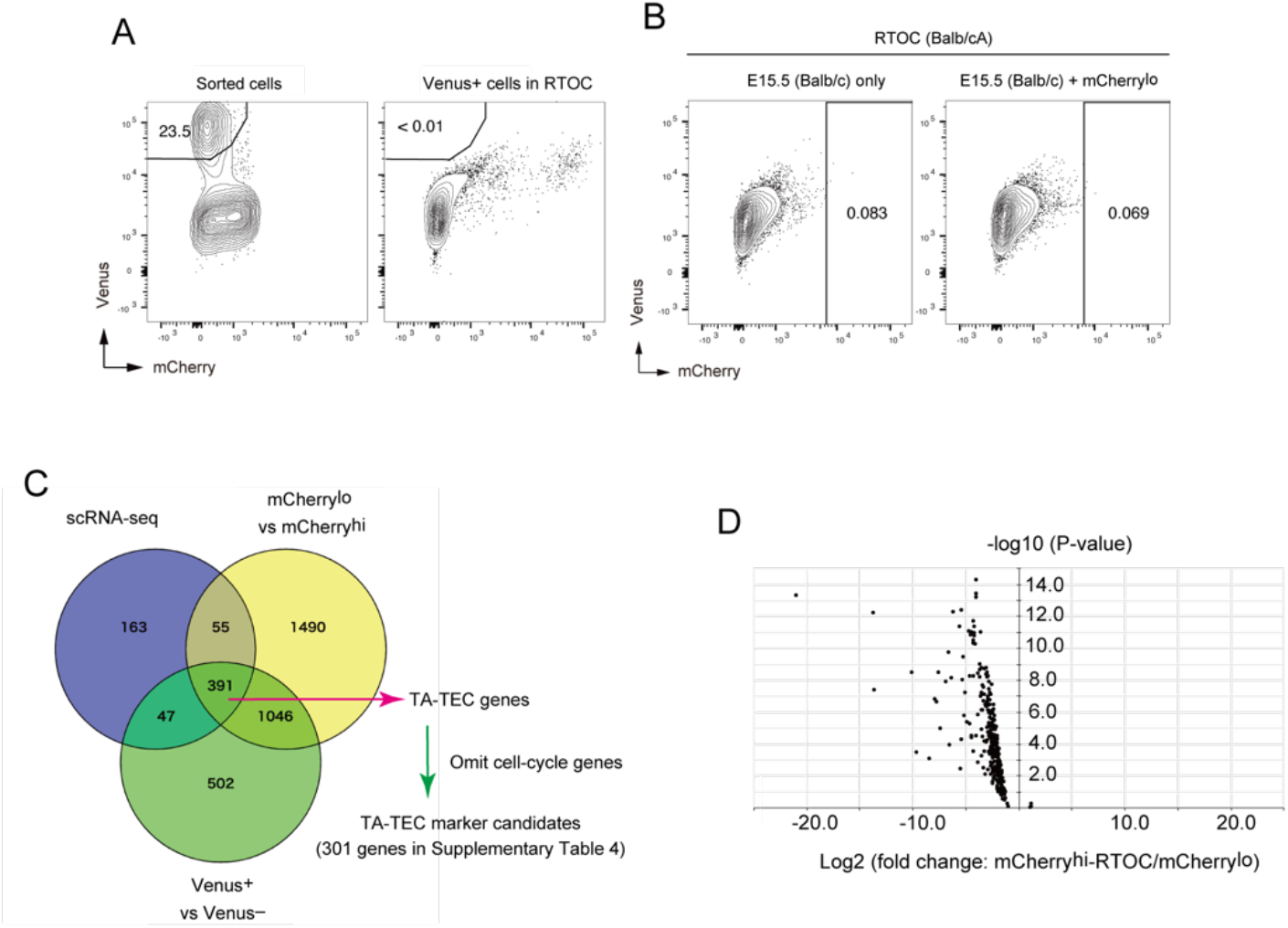
**A.** Ratio of Venus^+^ cells in sorted mCherry^10^ and RTOC **B.** Flow cytometric analysis of RTOC using allogenic fetal thymus (Balb/cA) **C.** Venn diagram of gene lists expressed in proliferating TECs at higher level in the 3 different RNA-seq datasets (mCherry^10^ vs mCherry^hl^ in Fig. 6, Venus+ vs Venus– in Fig. 4 and cluster R1 in Fig. 2). TA-TEC gene candidates were selected from the Venn diagram. TA-TEC marker gene candidates were selected by omitting cell cycle-related genes (GO:0007049 and Tirosh et al.^3^) form the TA-TEC gene candidates. The list of genes is summarized in Supplementary Table 4. **D**. Volcano plot for TA-TEC marker candidate expression in mCherry^10^ and mCherry^hi^ in RTOC.

**Figure 6-figure supplement 2.**
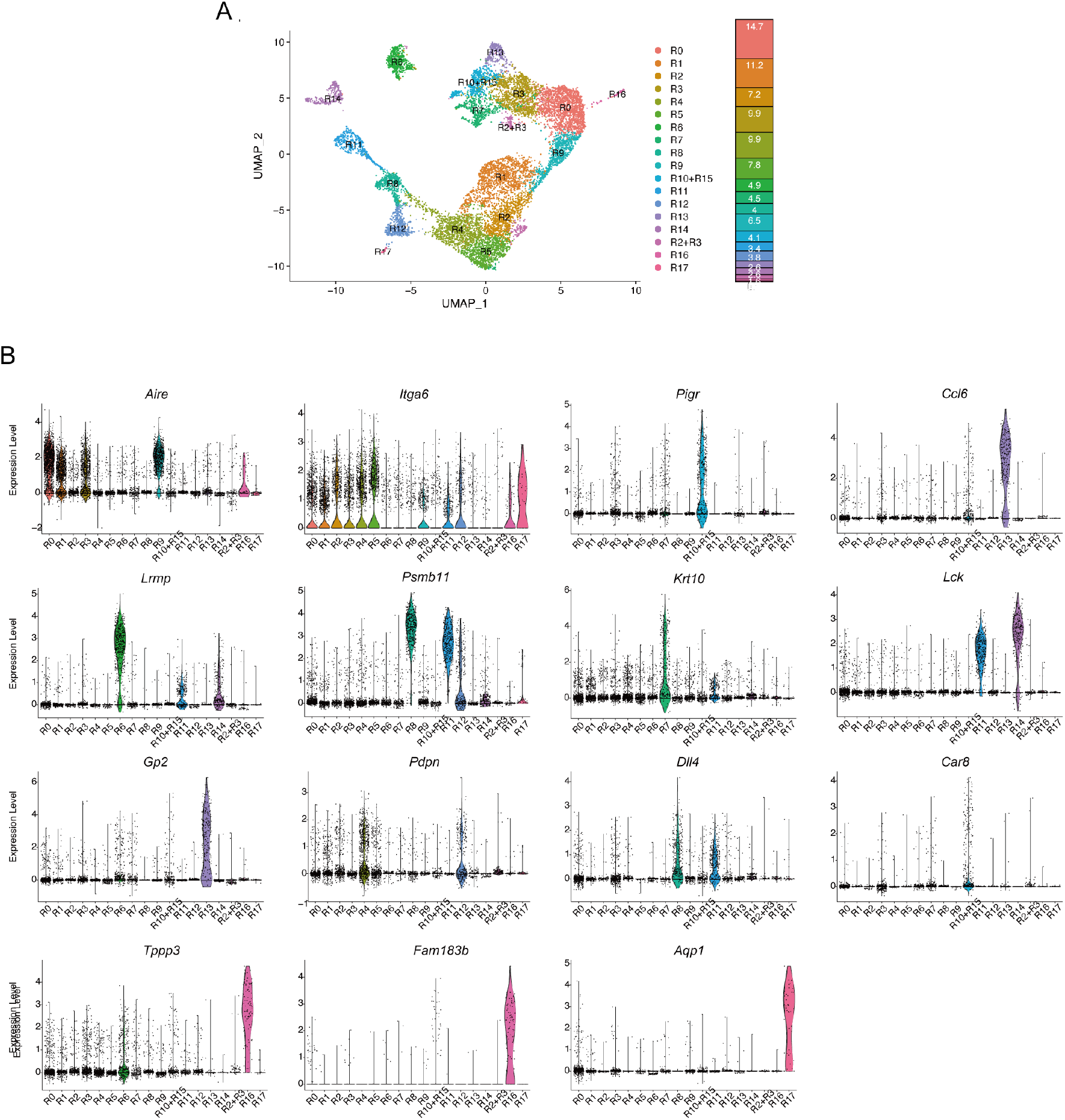
**A.** UMAP plot of droplet-based scRNA-seq and well-based scRamDA-seq data after their integration. Cell clusters are separated by colors and numbers in the plot. The graph on the right shows the percentages of each cluster in the total number of cells detected. Each cluster was assigned based on gene expression profile and corresponded with clusters in Fig. 2. **B**. Violin plots depicting expression level of typical TEC marker genes in each cluster.

**Figure 7-figure supplement 1.**
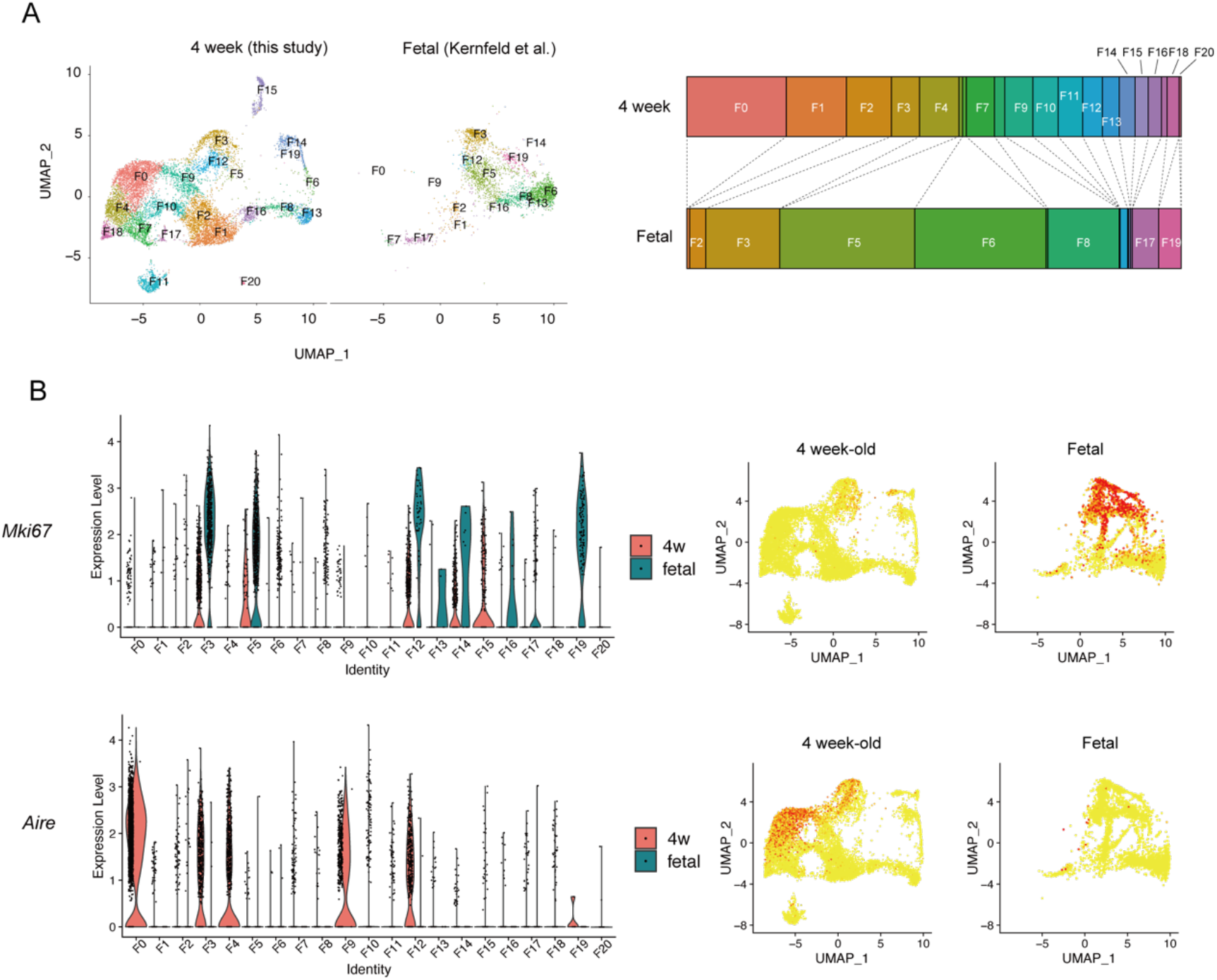
**A**. scRNA-seq data in this study (4-week-old mice) were integrated with scRNA-seq data reported by others^4^ **B.** Expression levels of *Mki67* (upper panels) and *Aire* (lower panels) a in the sub-cluster are shown in violin plots (left) and dot plots (right).

